# A single-cell transcriptomic atlas of the adult *Drosophila* ventral nerve cord

**DOI:** 10.1101/2019.12.20.883884

**Authors:** Aaron M. Allen, Megan C. Neville, Sebastian Birtles, Vincent Croset, Christoph D. Treiber, Scott Waddell, Stephen F. Goodwin

**Author notes:** For correspondence (A.M.A); (S.F.G). These authors contributed equally to this work (Aaron M. Allen, Megan C. Neville).

## Abstract

The *Drosophila* ventral nerve cord (VNC) receives and processes descending signals from the brain to produce a variety of coordinated locomotor outputs. It also integrates sensory information from the periphery and sends ascending signals to the brain. We used single-cell transcriptomics to generate an unbiased classification of cellular diversity in the VNC of five-day old adult flies. We produced an atlas of 26,000 high-quality cells, representing more than 100 transcriptionally distinct cell types. The predominant gene signatures defining neuronal cell types reflect shared developmental histories based on the neuroblast from which cells were derived, as well as their birth order. Cells could also be assigned to specific neuromeres using adult Hox gene expression. This single-cell transcriptional atlas of the adult fly VNC will be a valuable resource for future studies of neurodevelopment and behavior.

## Introduction

The adult *Drosophila* central nervous system (CNS) consists of the brain in the head capsule and the ventral nerve cord (VNC; also known as ventral nervous system) in the thorax (Court et al., 2017; Ito et al., 2014). The VNC receives and integrates sensory input from the periphery and sends this information to the brain in ascending neurons through the cervical connective (Tsubouchi et al., 2017). The brain, in turn, sends sensory-motor signals to the VNC via descending neurons (Namiki et al., 2018). The VNC transforms these signals into locomotor actions (Harris et al., 2015). It controls muscles in the thorax in unique ways, depending on whether it is steering the wings during flight or generating acoustic communication signals during both reproductive and agonistic behaviors (Clyne and Miesenbock, 2008; Jonsson et al., 2011; Shirangi et al., 2013; von Philipsborn et al., 2011). It coordinates muscles in the legs to walk, jump, groom, reach, touch, and taste (Bidaye et al., 2014; Card and Dickinson, 2008; Chen et al., 2018; Gowda et al., 2018; Harris et al., 2015; Howard et al., 2019; Kim et al., 2017; Mamiya et al., 2018; Mendes et al., 2013; Mendes et al., 2014; Seeds et al., 2014; Tuthill and Wilson, 2016; Wosnitza et al., 2013). The VNC also controls musculature in the abdomen relevant to copulation and reproduction including abdominal bending, attachment, and ejaculation in the male (Crickmore and Vosshall, 2013; Jois et al., 2018; Pan et al., 2011; Pavlou et al., 2016; Tayler et al., 2012), and sperm storage and oviposition in the female (Kimura et al., 2015; Lee et al., 2015).

These different functions are orchestrated by anatomically discrete segments of the VNC. Whereas the thoracic neuromeres control the legs and wings, the abdominal neuromere controls the abdominal muscles, gastric system, and reproductive organs. Much of the developmental mechanisms that generate and assemble the VNC have been characterized (Venkatasubramanian and Mann, 2019). Like other holometabolous insects, *Drosophila* undergo two stages of neurogenesis building two distinct nervous systems. Embryonic neurogenesis gives rise to the larval nervous system, and post-embryonic neurogenesis produces adult specific neurons. The structure of the adult nervous system is established in the embryo where pioneering neurons setup networks of tracks, which later developing neurons follow (Hartenstein, 2018). The adult VNC is comprised of 4 primary neuromeres, three thoracic (one per each pair of legs: the prothoracic neuromere, ProNm; mesothoracic neuromere, MesoNm and metathoracic neuromere, MetaNm) and one fused abdominal neuromere (ANm) (Court et al., 2017; Niven et al., 2008). The neuromeres, composed of one or more fused neuropils, are segment-specific parts of the CNS which process sensory signals and control movements of their specific segments. The thoracic neuromeres are homologous structures and are thus morphologically similar to each other (Smarandache-Wellmann, 2016). The neurons in these segments are derived from a set of repeating type I neuroblasts. Cells derived from a given neuroblast produce a lineage, which can be split into Notch^+^ and Notch^-^ hemilineages (Truman et al., 2010). The formation of tagmatic boundaries, which group these segments into morphological units along the body axis, is established through differential expression of Hox family transcription factors (TFs) (Angelini and Kaufman, 2005). Cell types within each neuromere are genetically encoded by developmental programs. However, it is unclear whether the mature terminal identity of an individual neuron can be determined from its adult transcriptome.

Although a large body of work has investigated how the adult *Drosophila* CNS is established from that of the embryo, many outstanding questions remain. It is for example unknown whether the TFs involved in establishing the cellular diversity of the nervous system also play a role in the form and function of these same neurons in the adult. Although certain features of individual neurons can be plastic, the identity of a terminally differentiated neuron is likely to remain stable throughout the animal’s life. It is therefore important to understand gene expression and regulation that permit mature neurons to preserve their subtype identity, morphology and connectivity and maintain neural circuit function throughout adulthood. The VNC of the genetically tractable vinegar fly, *Drosophila melanogaster*, is ideally suited for these studies.

Here we used single-cell RNA-sequencing to characterize the transcriptomes of individual cells in a 5-day old adult *Drosophila* VNC. We generated an atlas of the VNC with 26,768 single-cell transcriptional profiles that define more than 100 cell types. This analysis reveals that the VNC has a roughly equal number of inhibitory GABAergic neurons and excitatory cholinergic neurons, and that genes encoding preproneuropeptides are amongst the most highly expressed. The segmentally repeating nature and developmental history of the VNC is born out in the cellular transcriptomes, as maintained expression of Hox and several other neuronal lineage marker genes persist. In addition, these profiles provide many novel markers for known and new cell types. This single-cell atlas of the adult *Drosophila* VNC provides a useful resource to help connect cellular identity to behaviorally-relevant neural circuit function.

## Results

### Single-cell transcriptomic atlas of the adult VNC

We created a single-cell transcriptome atlas of the VNC using single-cell RNA-sequencing with 10x Chromium chemistry. We processed 80 VNCs, 20 per independent replicate (Figure 1A; see methods). Pooling the data, we recovered 26,768 high quality cells with a median of 1,170 genes and 2,497 transcripts per cell (Figure 1B). Lineage based counts of the adult specific post-embryonic neurons in the VNC range from 10-11,000 cells (Birkholz et al., 2015; Lacin et al., 2019), with a total number of cells in the VNC forecasted to be ∼16,000 (Lacin et al., 2019). Connectomic analyses of the adult VNC using Electron microscopy estimated a larger total number of ∼20,000 cells (Bates et al., 2019). Based on these prior numbers, our cell atlas has 1.3-1.7x coverage of the VNC. We performed a Canonical-Correlation Analysis (CCA) on the transcriptomes and reduced the first 45 dimensions into two t-SNE dimensions (van der Maaten, 2014) resulting in 120 distinct clusters (Figure 1B, Figure1-Figure supplement 1). We evaluated the clustering resolution with a clustering tree, comparing the relationship between clusters at different resolutions (Figure 1-Figure supplement 2). A cluster resolution of 12 was chosen as it best resolved established substructures within our data (as described below). Cells from independent experimental replicates were equally distributed across the clusters with an alignment metric of 94.6% (Figure 1-Figure supplement 3A). In addition, a similar number of expressed genes (nGene) and transcripts (nUMI) were retrieved in each replicate (Figure 1-Figure supplement 3B), and the number of genes and transcripts showed a uniform distribution across the t-SNE (Figure 1-Figure supplement 3C). Lastly, we compared the expression levels observed in this single-cell data set with those reported from bulk sequencing, to evaluate the extent to which the tissue dissociation and the 10x procedure effect gene expression. The filtered single-cell data showed high correlation to FlyAtlas’ bulk VNC sequencing data (Figure 1-Figure supplement 4) (Leader et al., 2018).

**Figure 1:**
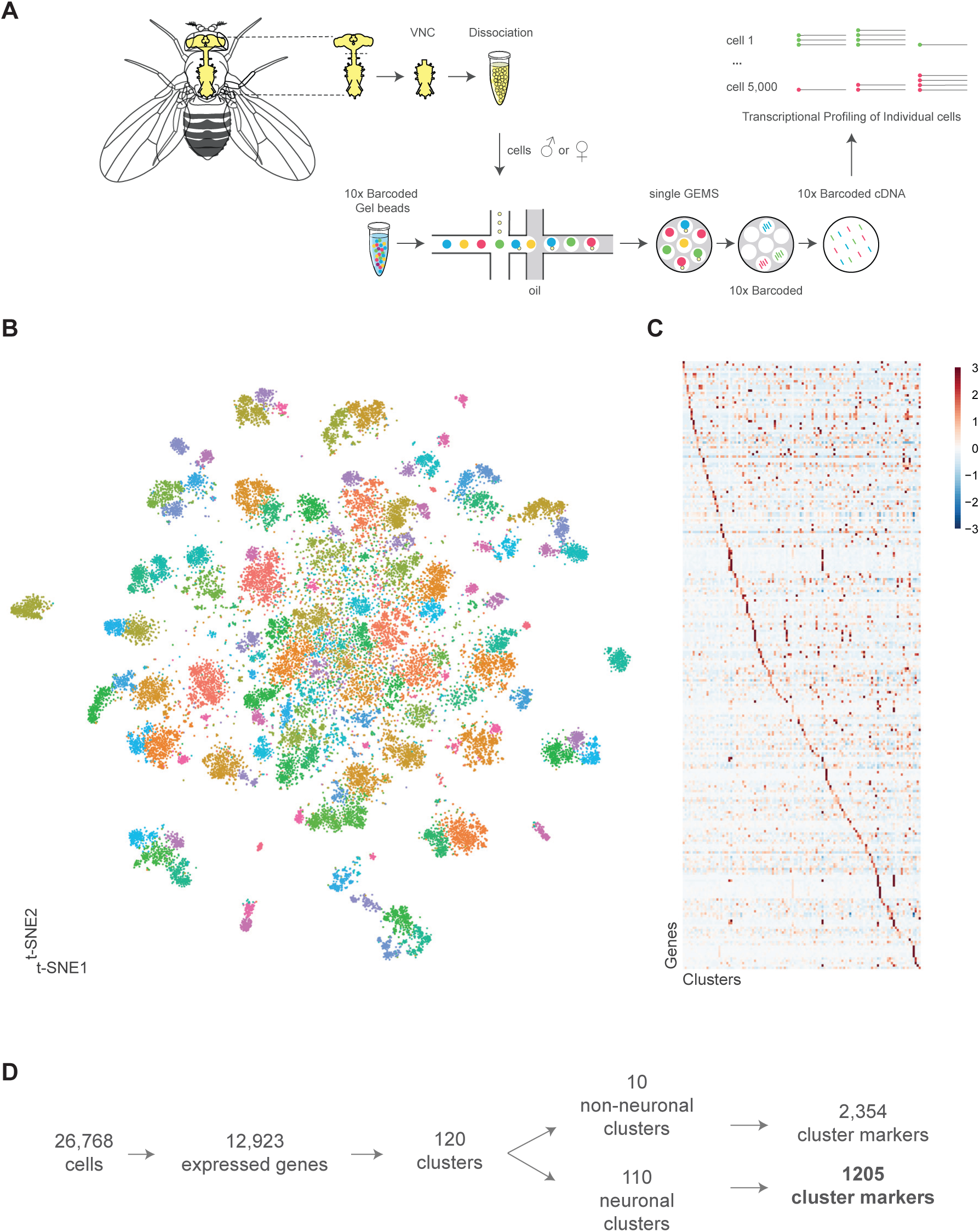
Single cell sequencing of 5-day old adult Drosophila VNC. A. Schematic workflow showing single cell sequencing data generation. Male and female Drosophila VNCs were dissected and dissociated prior to droplet encapsulation of individual cells with barcoded beads, forming gel beads in emulsion (GEMs). Following barcode incorporation, molecular amplification and sequencing the transcriptional profiles of individual cells were determined. B. Two-dimensional representation (t-SNE) of 26,768 Drosophila VNC cells grouped into 120 clusters. Clusters were assigned by shared nearest neighbor, using 45 CCA dimensions. Each dot is a cell colored by cluster identity. C. Heatmap showing scaled, log-normalized expression of top 5 cluster-discriminative genes per cluster. D. Flow diagram representing neuronal and non-neuronal cluster identification, including total number of genes identified as cluster markers.

### Cluster-defining marker genes

The 120 tSNE clusters are defined by unique combinations of significantly enriched genes that we refer to as cluster markers (Figure 1C, Figure1-Figure supplement 1, Figure 1-source data 1). These cluster markers are likely to be important for the development and/or maintenance of the cell types represented by the clusters. We defined 110 neuronal clusters using expression of the known neuron-specific genes, *embryonic lethal abnormal visio*n (*elav*), *neuronal Synaptobrevin* (*nSyb*) and the long non-coding (lnc) RNA *noe* (*noe*) (Davie et al., 2018; DiAntonio et al., 1993; Kim et al., 1998; Robinow et al., 1988). These neuronal clusters can be distinguished from each other by differential expression of 1205 additional cluster markers (Figure 1D, Figure 1-source data 1). Functional enrichment analysis of these cluster-defining marker genes identified those encoding immunoglobulin (Ig)-like domains as most significantly enriched (Figure 2A), including many cell adhesion molecules that are known to establish neuronal connectivity during development (Özkan et al., 2013). Specifically, many members of the Immunoglobulin superfamily (IgSF) continue to be expressed in the adult and are significant cluster markers (Figure 2B; Figure 2-Figure supplement 1A). The defective proboscis extension response (Dpr) subfamily, and their binding partners, the Dpr-interacting protein (DIP) subfamily, are markers for over 50% of neuronal clusters identified in the adult VNC (Figure 2-Figure supplement 1B). Dpr and DIP genes provide a complex interaction network regulating neural circuit assembly and have been proposed to act as neuronal ‘identification tags’ during development. Distinct combinations of Dpr and DIP proteins are expressed by different neuronal classes in the developing optic lobe, the antennal lobe, and the VNC (Carrillo et al., 2015; Özkan et al., 2013). Specific combinations of Beat and Side IgSF proteins play an important role in the sculpting the nervous system (Özkan et al., 2013; Pipes et al., 2001). Almost all Beat and Side family members were found as cluster markers in the VNC (Figure 2B), and their expression was largely mutually exclusive, with 80% of the identified clusters enriched for either Beats or Sides (Figure 2-Figure supplement 1C). The enrichment of IgSFs in the adult VNC supports their importance in establishing cell type specificity and suggests an ongoing role in maintenance of neuronal connectivity in the mature nervous system (Figure 2B).

**Figure 2:**
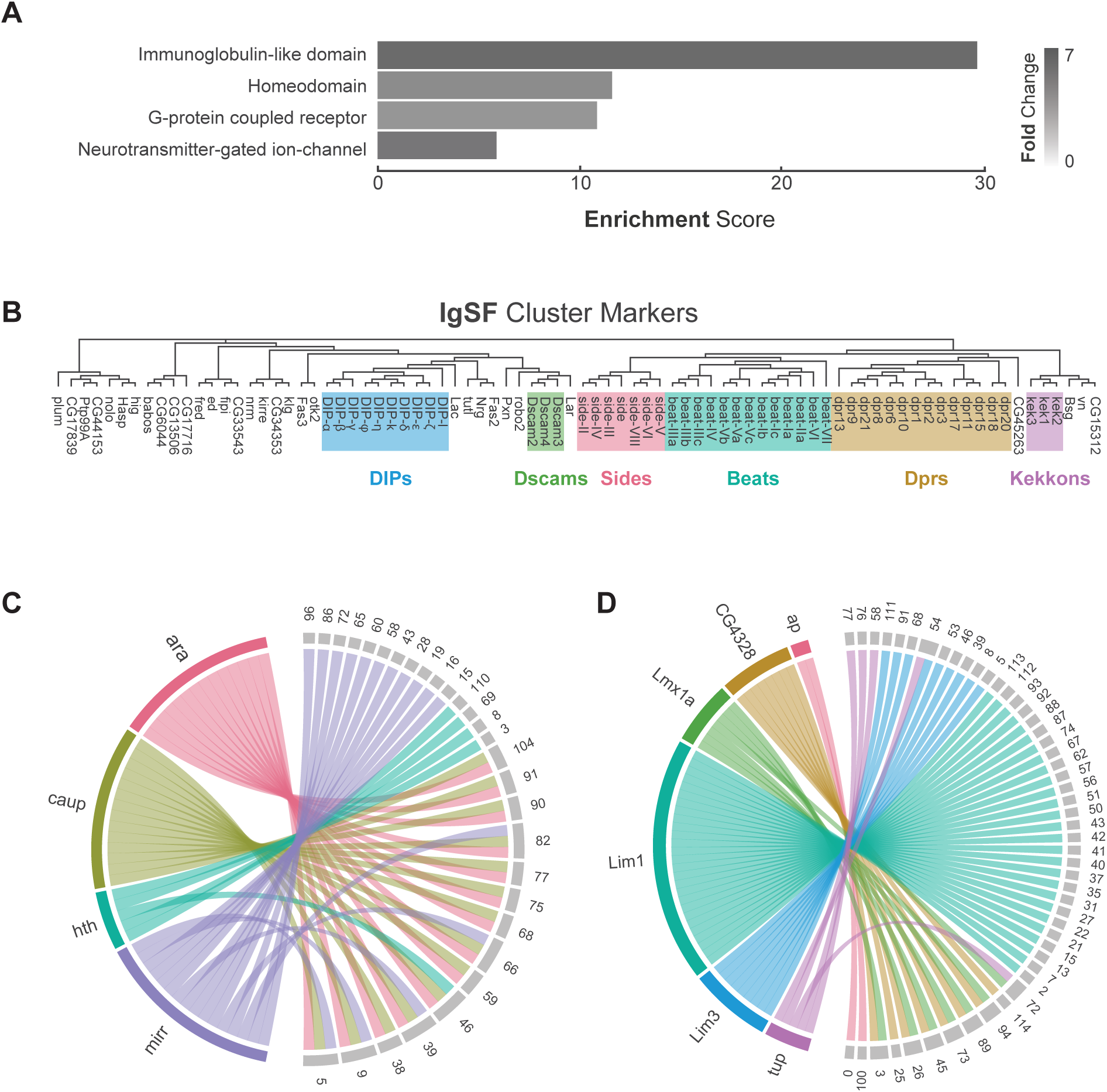
Characterization of neuronal cluster markers. A. Functional analysis of neuronal cluster markers (DAVID) showing representative functional terms (>5 fold-change) and their corresponding enrichment scores (representing mean p-values). B. Phylogenetic tree of IgSF neuronal cluster markers. IgSF subfamily members are highlighted in color. C. Chord diagram showing Tale Homeobox TF cluster markers (left, in color) with the clusters for which they are significantly enriched (right, in grey). D. Chord diagram showing LIM Homeobox TF cluster markers (left, in color) with the clusters for which they are significantly enriched (right, in grey). B-D. Gene were classified based on Flybase Gene Group annotations (www.flybase.org).

G-protein coupled receptors (GPCRs) were also significantly over-represented as cluster markers (Figure 2A, Figure 2-Figure supplement 2A). GPCRs play a critical role in intercellular communication by interacting with a diverse group of extracellular ligands such as neurotransmitters, modulatory neuropeptides and biogenic amines, peptide hormones, and gases (Hanlon and Andrew, 2015). We observed a median of 5 GPCRs expressed per cell and a median of 22 transmembrane receptors in total per cell. Although many GPRCs are significantly enriched in the individual clusters, they were, in general, expressed at low levels, in few cells per cluster, and across many clusters (Figure 2-Figure supplement 2A). The neuropeptide receptor family members had more inter-cluster variation in expression levels than the other GPCR subtypes, suggesting that they confer an added level of specialization. We also saw that 80% of biogenic amine receptor family members, a diverse group of neurotransmitter receptors sensitive to biogenic amine neurotransmitters including dopamine, octopamine, serotonin and tyramine, are found as cluster markers (Figure 2-Figure supplement 2B). Odorant receptors, gustatory receptors, ionotropic receptors, and pickpocket sodium channels are predominantly found in first-order sensory neurons (Joseph and Carlson, 2015) and in general, were not expressed in the VNC, as expected. There were exceptions, however, as *Or63a*, *Gr28b*, *Ir76a*, *ppk31* are all expressed in the VNC, albeit at low levels and in few cells. Neurotransmitter-gated ion-channels were also highly enriched as neuronal cluster markers (Figure 2A, Figure 2-Figure supplement 3, Figure 1-source data 1), including 8 of the 10 nicotinic acetylcholine receptor subunits (Littleton and Ganetzky, 2000). Expression of transmembrane receptors that provide fast- and slow-acting responses to neurotransmitters, therefore, defines cellular identify.

Finally, over 20% of all *Drosophila* TFs were cluster markers in the adult VNC, the most enriched class of which was the homeodomain (HD) family (Figure 2A, Figure 2-Figure supplement 4A). Homeodomain TFs play central roles in establishing regional-, tissue- and cell-specific fates (Bürglin and Affolter, 2016). We observed a median of 5 HD TFs per cell, and a median of 52 TFs in total per cell. We detected high variance in HD TF expression levels across clusters (Figure 2-Figure supplement 4B). Four members of the three amino acid loop extension (TALE) class of HD TFs mark specific clusters. This included all the three members of the evolutionarily conserved *Iroquois* gene complex (Iro-C) *araucan* (*ara*), *caupolican* (*caup*), and *mirror* (*mirr*) (Cavodeassi et al., 2001). The Iro-C arose through two duplication events, one ancient event in arthropods led to independent *ara*/*caup* and *mirr* genes, followed by a more recent event in dipterans which gave rise to the *caup* and *ara* genes (Kerner et al., 2009). Cluster expression of Iro-C genes appears to reflect this evolutionary history. Whereas *caup* and *ara* overlap as cluster markers, *mirr* expression is only partially overlapping and *mirr* is an independent marker for 11 additional clusters, suggesting specialization (Figure 2C). In addition, we found largely mutually exclusive enrichment of individual members of the LIM class of HD TFs defining cell clusters in the VNC (Figure 2D). LIM TFs are known to specify distinct neuronal identities in the embryo (Thor et al., 1999). In the VNC only *Lmx1a* and *CG4328* appear to be co-expressed which given their genomic linkage likely represents a relatively recent tandem duplication event. These findings suggest that aspects of the TF-code that establishes cellular identity during development actively maintains neuronal identity throughout the life of the animal (Deneris and Hobert, 2014).

### Hox genes define neuromere identity

Hox genes encode homeodomain proteins that confer positional identities along the antero-posterior axis in all bilaterian animals (McGinnis and Krumlauf, 1992). In both vertebrates and invertebrates, the Hox family of transcription factors are known to govern key aspects of nervous system development, notably the formation of neuromuscular networks (Philippidou and Dasen, 2013). The neuromeres of the VNC show distinct segment-specific properties under Hox gene control (Baek et al., 2013; Suska et al., 2011).

The Hox genes *Antennapedia* (*Antp*), *Ultrabithorax* (*Ubx*), *abdominal-A* (*abd-A*), and *Abdominal-B* (*Abd-B*), which specify the thoracic, and abdominal segments of the fly VNC, define specific clusters in the single-cell atlas (Figure 3A). Hox gene expression patterns were significantly anti-correlated at the individual cell level, except for *abd-A* and *Abd-B* (Figure 3B). *Antp* and *Ubx* are both expressed throughout the t-SNE plot, with many clusters expressing both, albeit in mutually exclusive cells, while *abd-A* and *Abd-B* expression is more restricted and overlapping (Figure 3C). Immunostaining the adult VNC with antibodies against these Hox proteins showed belts of expression along the anterior-posterior axis (Figure 3D). These patterns of expression are different to those in the larva but are consistent with those observed in the mid-pupal stage (Baek et al., 2013). Antp protein is most highly expressed in the mesothoracic neuromere (MesoNm), whereas Ubx expression is highest in the metathoracic neuromere (MetaNm). Abd-A and Abd-B protein showed overlapping expression restricted to the abdominal neuromere (ANm), with Abd-A expressed more anteriorly than Abd-B. By combining the neuromere specific (or enriched) expression revealed by antibody staining, with the high level of anti-correlation in the single-cell data, we can separate the single-cell clusters into approximations for each neuromere (Figure 3E).

**Figure 3:**
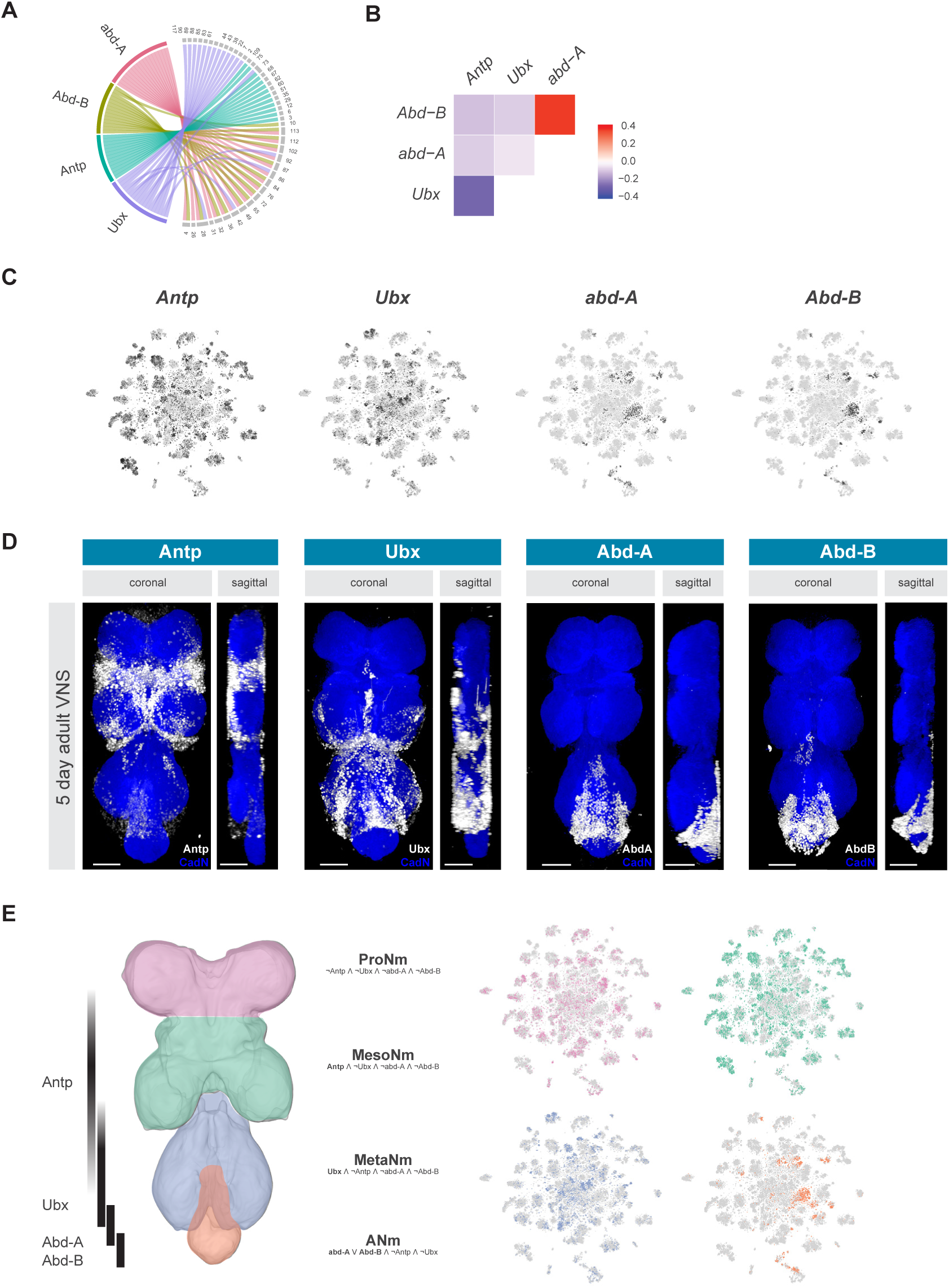
Neuromeres are defined by Hox gene expression in the adult VNC. A. Chord diagram showing Hox genes found as cluster markers (left, in color) with the clusters for which they are significantly enriched (right, in grey). B. Heatmap of Pearson correlation coefficient for Hox gene expression. C. t-SNE plot of Hox genes. Expression shown in black, intensity is proportional to the log-normalized expression levels. D. Visualisation of Hox protein expression in the 5-day old adult VNC. Coronal and sagittal views of anti-Antp, -Ubx, -Abd-A, and -Abd-B (white). Neuropil counterstained with anti-Cad-N (blue). Scale bars = 50 μm. E. Schematic representing bands of Hox expression along the anterior-posterior axis of the VNC (left). Cells assigned to neuromeres in VNC t-SNE plots (right) based on differential Hox gene expression: ProNm (pink), MesoNm (green), MetaNm (blue) and ANm (orange). Genes included in defining each neuromere in bold.

### Neuroblast lineage identity

Each neuromere contains a repeating set of neuroblast lineages. We therefore used prior knowledge of biomarkers defining the development of post-embryonic lineages (Bossing et al., 1996; Schmid et al., 1999) to further annotate the 5-day old adult VNC single-cell atlas. All known lineage biomarkers showed continued expression in the adult VNC, and most were found as significantly enriched cluster markers (Figure 4A and B; Figure 4-Figure supplement 1; Figure 1-source data 1). These extant markers therefore allowed us to assign potential hemilineage identities to many of the cell clusters (Figure 4C). We also used fast-acting neurotransmitter (FAN) identity as an additional marker to specify hemilineages, since FAN usage is acquired in a lineage-dependent manner in post-embryonic neurons in the VNC (Lacin et al., 2019). For example, cluster 54 has enriched expression of lineage markers *Lim3*, *tailup* (*tup), abnormal chemosensory jump 6* (*acj6*), and the glutamatergic neuron marker *Vesicular glutamate transporter* (*VGlut*), which collectively identifies this cluster as hemilineage 9B. The correspondence between cluster markers and hemilineage markers shows that cell cluster identity is closely linked to hemilineage identity. We observed a strong concordance between the number of cells within our predicted hemilineages and previous cell counts (Figure 4-Figure supplement 2A) (Birkholz et al., 2015; Lacin et al., 2019). This correlation of cell counts further supports our atlas having an approximately 1.5x coverage of the VNC.

**Figure 4:**
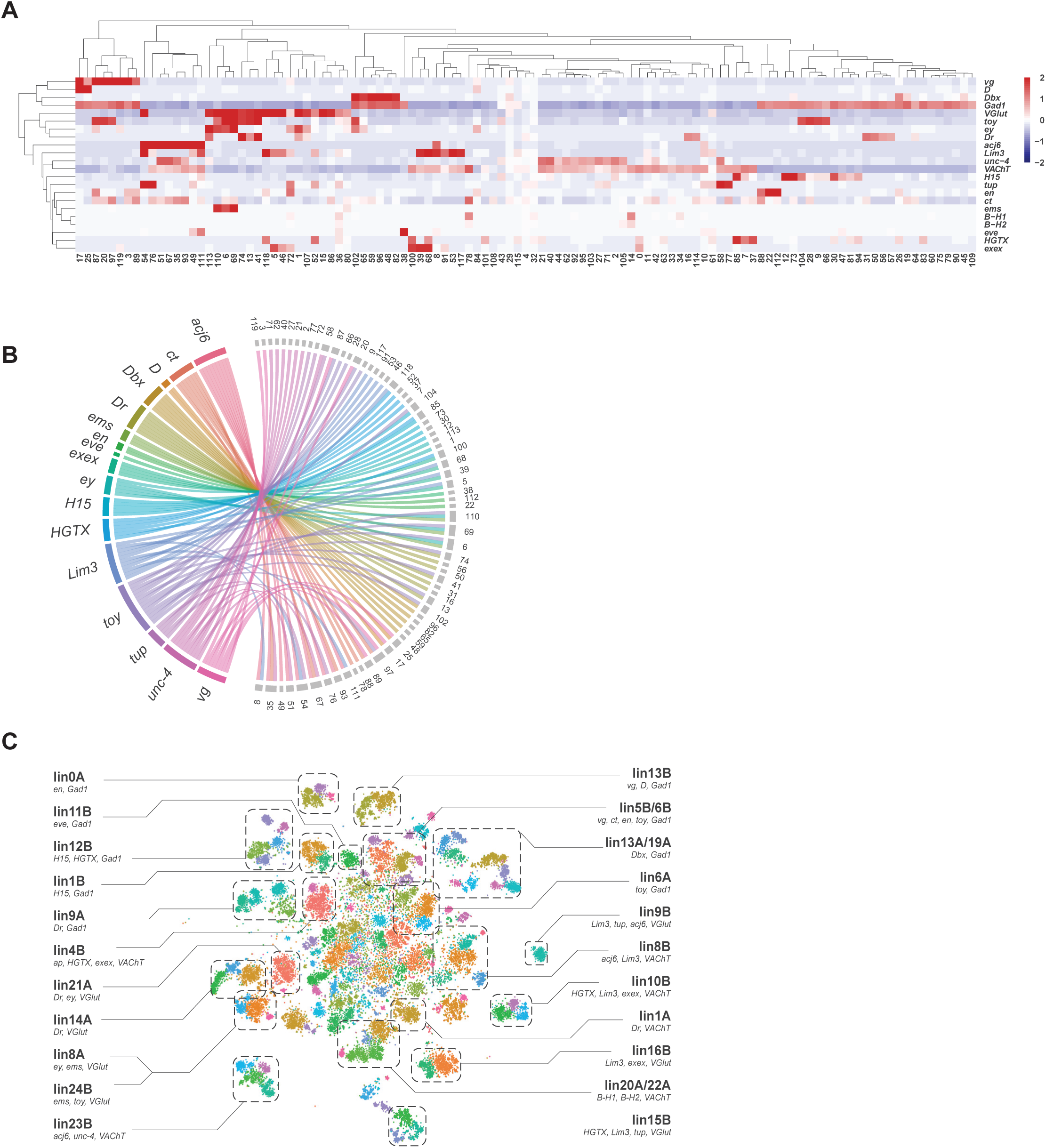
Cellular identities defined by developmental lineages. A. Heatmap of the mean scaled log-normalized expression, by cluster (bottom), of the neuroblast hemilineage and FA-neurotransmitter markers (right). Dendograms represent hierarchical clustering. B. Chord diagram showing neuroblast lineage markers (left, in color) with their associated clusters for which there are significantly enriched (right, in grey). C. t-SNE plot of neuronal cells with clusters predicted to represent defined lineages based on the expression of lineage marker genes (shown below lineage names).

Many of the established lineage markers are expressed at low levels and in few cells (e.g., *B-H1/2*, *exex*, *tup*), while others are robustly expressed (e.g., *acj6*, *toy*, *Lim3*) (Figure 4-Figure supplement 1). Nevertheless, the restricted cluster-specific expression of these established markers and co-expression with more robust novel markers allowed us to assign cells to several hemilineages. At present some hemilineages cannot be defined with certainty due to the lack of specific sets of markers. For instance, *unc-4* is currently the lone marker for many cholinergic hemilineages, including 7B, 12A, 18B and 19B. More work will be needed to resolve the multiple cholinergic clusters in the center of the t-SNE that express *unc-4* and are therefore likely to represent these hemilineages (Figure 4-supplement 1). Conversely, clusters 6, 69, and 110 express markers for both hemilineages 8A (*ems*, *ey*, *VGlut*) and 24B (*ems*, *toy*, *VGlut*), suggesting that these lineages exhibit very similar transcriptional profiles and resolving them may require more cells (Figure 4C).

Many predicted hemilineages contain multiple clusters (Figure 4C). We propose that these clusters represent distinct variations within a lineage, partly informed by birth order and partly related to thoracic vs. abdominal anatomical position. Consistent with a subdivision based on birth order, many of our predicted hemilineages contain a distinct subgroup of *broad* (*br*) expressing cells (Figure 4-Figure supplement 2B). A specific isoform of *br* continues to be expressed in the adult and marks cells born around L2-L3 ecdysis (Zhou et al., 2009). Other lineage sub-division was evident when visualizing expression of neurodevelopmental genes such as *prospero* (*pros*), *datilografo* (*dati*), *maternal gene required for meiosis* (*mamo*), and *IGF-II mRNA-binding protein* (*Imp*) (Figure 4-Figure supplement 2C). Expression of *pros* and *Imp* in the VNC were negatively correlated (Figure 4-Figure supplement 2D), as previously observed in the adult brain (Davie et al., 2018). In contrast, expression of *pros* was highly correlated with that of *dati*, while *Imp* and *mamo* were highly correlated, suggesting these pairs of genes may be co-regulated in the adult (Figure 4-Figure supplement 2D). *pros* and *Imp* also showed an anatomical bias, with *pros* being negatively correlated with abdominally expressed *abd-A* and *Abd-B*, while *Imp* is positively correlated with these markers (Figure 4-Figure supplement 2E). However, we did not observe a striking physiological bias, as neither *pros* or *Imp* showed correlation with fast-acting neurotransmitter markers (Figure 4-Figure supplement 2F). It has previously been shown that *pros* labels late-born motor neurons from lineage 15B (Baek et al., 2013), and that a trade-off of *Imp* and *pros* expression is essential for cell cycle exit of post-embryonic type I neuroblasts in the VNC (Maurange et al., 2008; Yang et al., 2017). Studies investigating the transcriptomes of mature (embryonic) vs. immature (post-embryonic) neurons in the larval VNC have shown that *Imp* is enriched in mature neurons, and *pros* is enriched in immature neurons (Etheredge, 2017). We propose that the continued expression of *Imp* in the adult VNC labels early-born and embryonic neurons while *pros* labels late-born post-embryonic neurons.

### Gene expression defines neuronal subtypes within a hemilineage

Hemilineage 23B marked by *acj6*, *unc-4*, and *VAChT* has four distinct clusters that can only be partially explained by potential birth order and neuromere identity (Figure 4C). *acj6* is also a marker for the cholinergic hemilineage 8B and the glutamatergic hemilineage 9B (Figure 4C) (Lacin et al., 2019). We examined *acj6* expressing cells in more detail by visualizing the anatomical distribution of *acj6* expressing cell bodies and neuronal projections in the VNC using *acj6^GAL4^* to express the GAL4-responsive dual reporter *UAS-Watermelon* (WM), which simultaneously marks the plasma membrane with GFP and the cell nucleus with mCherry (Figure 5A) (Lee et al., 2018). We also validated the accuracy of *acj6^GAL4^* using an anti-Acj6 antibody (Clyne et al., 1999; Figure 5-Figure supplement 1). Consistent with previous reports, *acj6^GAL4^* labeled distinct clusters of cells in the three thoracic neuromeres of the VNC, which predominantly innervate the leg neuropils. (Figure 5A) (Harris et al., 2015; Shepherd et al., 2019). The *acj6* expressing neurons were confirmed as hemilineages 8B, 9B, and 23B (D. Shepherd, pers. comm.) on the basis of their primary neurite projections (Shepherd et al., 2016).

**Figure 5:**
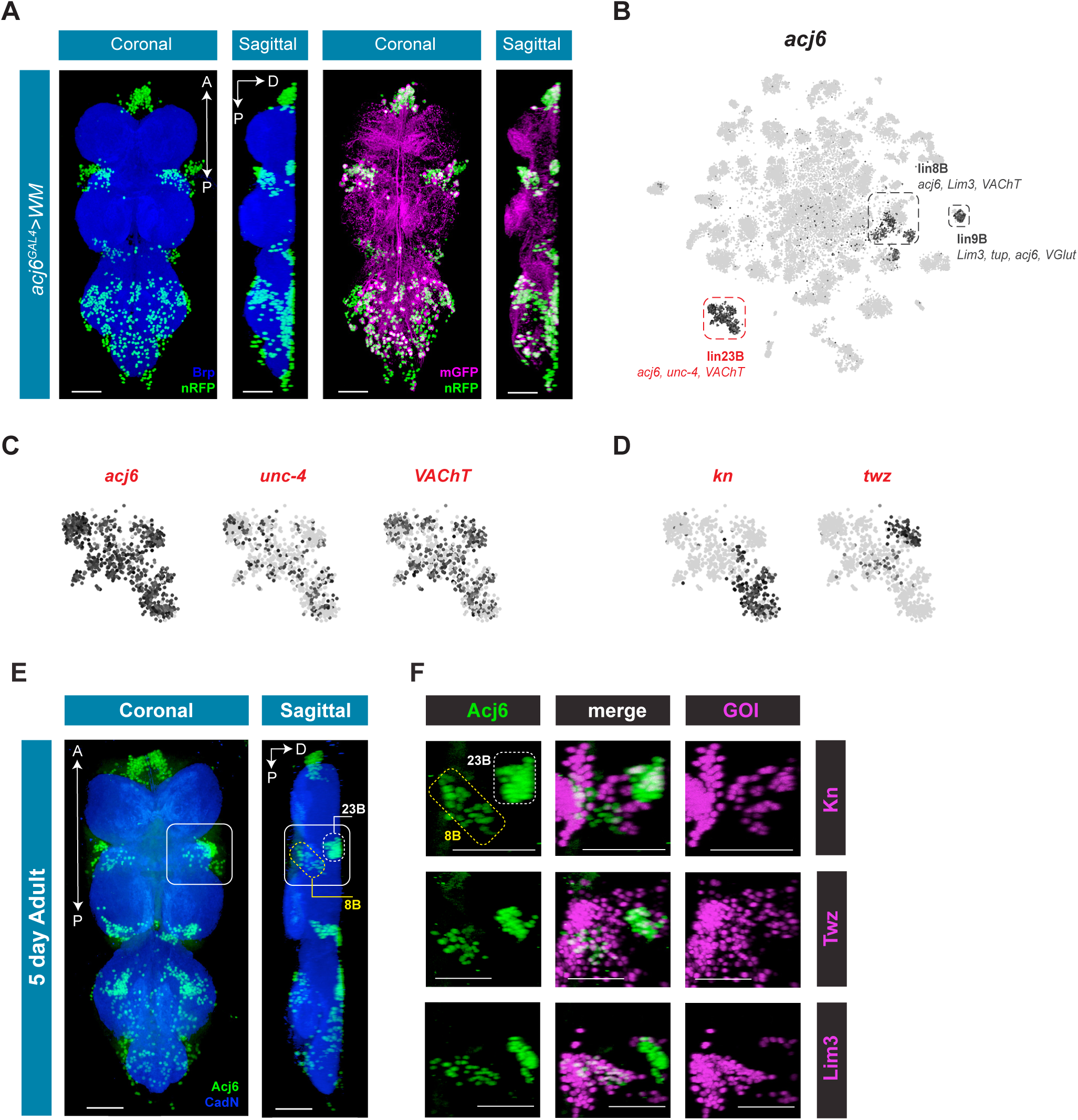
acj6 expression, hemilineage identity, and novel lineage marker co-expression in the VNC. A. Visualisation of acj6 expressing cells in the 5-day old adult VNS. Coronal and sagittal views of *acj6^GAL4^* driving expression of *UAS-WM*, enabling the visualization of cell nuclei (nRFP; green) and neuronal morphology (mGFP; magenta). Neuropil is counterstained (Brp; blue). A, Anterior, P, posterior, D, dorsal; scale bars = 50 μm. B. t-SNE plot of *acj6* expression in the VNC. Predicted hemilineage 8B, 9B, and 23B, based on known markers (shown below) are highlighted. C. t-SNE plots of predicted hemilineage 23B, showing expression of markers *acj6*, *unc-4*, and *VAChT*. D. t-SNE plots of predicted hemilineage 23B, showing expression of novel sub-type markers *kn* and *twz*. B-D. Expression shown in black, intensity is proportional to the log-normalized expression levels. E. Acj6 expression in the 5-day old adult VNC; coronal view (left) and sagittal view (right). Solid white box encompasses the posterior prothoracic/anterior mesothoracic region, containing cells of hemilineages 23B, dorsally (dashed white box), and 8B, ventrally (dashed yellow box). A, Anterior, P, posterior, D, dorsal; Neuropil is counterstained (CadN; blue) scale bars = 50 μm. F. Close-up sagittal views of posterior prothoracic/anterior mesothoracic region; Acj6 expression (anti-Acj6; green; left column) in hemi-lineage 23B cells (dashed white box) and 8B cells (dashed yellow box). Acj6 expression overlaps (merge, white cells) with *kn^GAL4^* and *twz^GAL4^* driven *UAS-Stinger* expression (nGFP; magenta; right column) in cells of hemilineage 23B (top two rows); and with *Lim3^GAL4^* driven *UAS-Stinger* expression (nGFP; magenta) in cells of hemilineage 8B (bottom row). GOI = gene of interest.

The *acj6* expressing clusters in the single-cell data set are highlighted in Figure 5B. Based on co-expression of additional markers, we could definitely assign some *acj6*-expressing hemilineages within the VNC data; lin8B expresses *acj6*, *Lim3*, and *VAChT*; lin9B *acj6*, *Lim3*, *VGlut*, and *tup*; Lin23B *acj6*, *unc-4*, and *VAChT* (Figure 5C). We next used novel markers identified in the single-cell data to determine whether these could account for the clusters within the 23B hemilineage. The *knot* (*kn/collier*) and *tiwaz* (*twz*) (Crozatier et al., 1996; Williams et al., 2014) genes are expressed in distinct subsets of hemilineage 23B (Figure 5D), whereas the established marker *Lim3* is co-expressed with *acj6* in hemilineages 8B and 9B (Figure 5B) (Lacin and Truman, 2016). We validated the co-expression evident in the single-cell data using GAL4 lines that represent *kn*, *twz*, and *Lim3* expression, and co-stained these VNCs with anti-Acj6 antibody (Figure 5E and F; Figure 5-Figure supplement 2). Hemilineage 23B is located dorsolaterally at the posterior edge of each of the three thoracic neuromeres, whereas 8B is ventrolateral at the anterior side of each thoracic neuromere (Figure 5E) (Lacin et al., 2019; Shepherd et al., 2019). Focusing on the ProNm-MesoNm border, which encompasses prothoracic 23B and mesothoracic 8B, we found that a subset of 23B co-expressed *kn* and a subset co-stained for *twz*, while *Lim3* expression was restricted to 8B (Figure 5F). Full expression patterns in the VNC are shown in Figure 5-Figure supplement 2. These findings highlight the predictive power of the single-cell transcriptome to identify new markers that refine our understanding of hemilineage subtypes which contribute to neuronal cell diversity in post-embryonic lineages.

### Classification using fast-acting neurotransmitters

A recent comprehensive map of fast-acting neurotransmitter (FAN) usage across the VNC found that neurons within a post-embryonic hemilineage use the same neurotransmitter; either acetylcholine, gamma-aminobutyric acid (GABA), or glutamate, unifying their shared developmental and functional identities (Lacin et al., 2019). We examined FAN usage in the VNC using expression of established biosynthetic and vesicular loading markers: *Choline acetyltransferase* (*ChAT*) and *Vesicular acetylcholine transporter* (*VAChT*) for acetylcholine, *VGlut* for glutamate, and *Glutamic acid decarboxylase 1* (*Gad1*) and *Vesicular GABA Transporter* (*VGAT*) for GABA (Figure 6A). Clusters showed mutually exclusive enrichment of *VAChT*, *VGlut*, and *Gad1* (Figure 6B). However, this exclusivity partially breaks down at the cell-by-cell level. Although expression levels of *VAChT*, *VGlut*, and *Gad1* were negatively correlated (Figure 6-Figure supplement 1A), we also observed significant levels of co-expression, with more than 28% of cells expressing at least two of these markers (Figure 6-Figure supplement 1B). A similar pattern of co-expression was observed in single-cell data from the adult brain (Croset et al., 2018; Davie et al., 2018). However, immunostaining of the intact adult VNC suggested that cytoplasmic co-expression of these markers does not occur (Lacin et al., 2019). Co-expression in the VNC may therefore represent a technical artifact of the single-cell approach. Since the cells cluster due to hemilineage identity and hemilineages are restricted to a single FAN, we assigned FAN identity by comparing the average expression of these FAN markers at the cluster level (Figure 6-Figure supplement 1C). Amongst the co-expressing cells, the expression of FAN makers is negatively correlated, and cells assigned to one FAN identity tended to show higher expression of the corresponding marker (Figure 6-Figure supplement 1D). With these criteria (see methods), we estimate that the VNC is 40% cholinergic, 38% GABAergic, and 18% glutamatergic. The remaining 4% of neurons did not show a strong signature for any FAN (Figure 6C and D). These proportions differ to those in the adult brain (Croset et al., 2018; Davie et al., 2018), with inhibitory GABAergic neurons being much more prominent in the VNC (38% vs 15% in the adult midbrain). The inhibitory glutamate receptor, *GluClα* (Cully et al., 1996), was broadly expressed in these data and is particularly enriched in cluster 103, suggesting that a portion of glutamatergic signaling is also inhibitory. We expect this relative abundance of inhibitory signaling reflects the importance of inhibition in the initiation, maintenance, and termination of complex motor programs (Burrows, 1992; Grillner, 2006), especially in coordination between bilateral and intersegmental circuits (Gowda et al., 2018).

**Figure 6:**
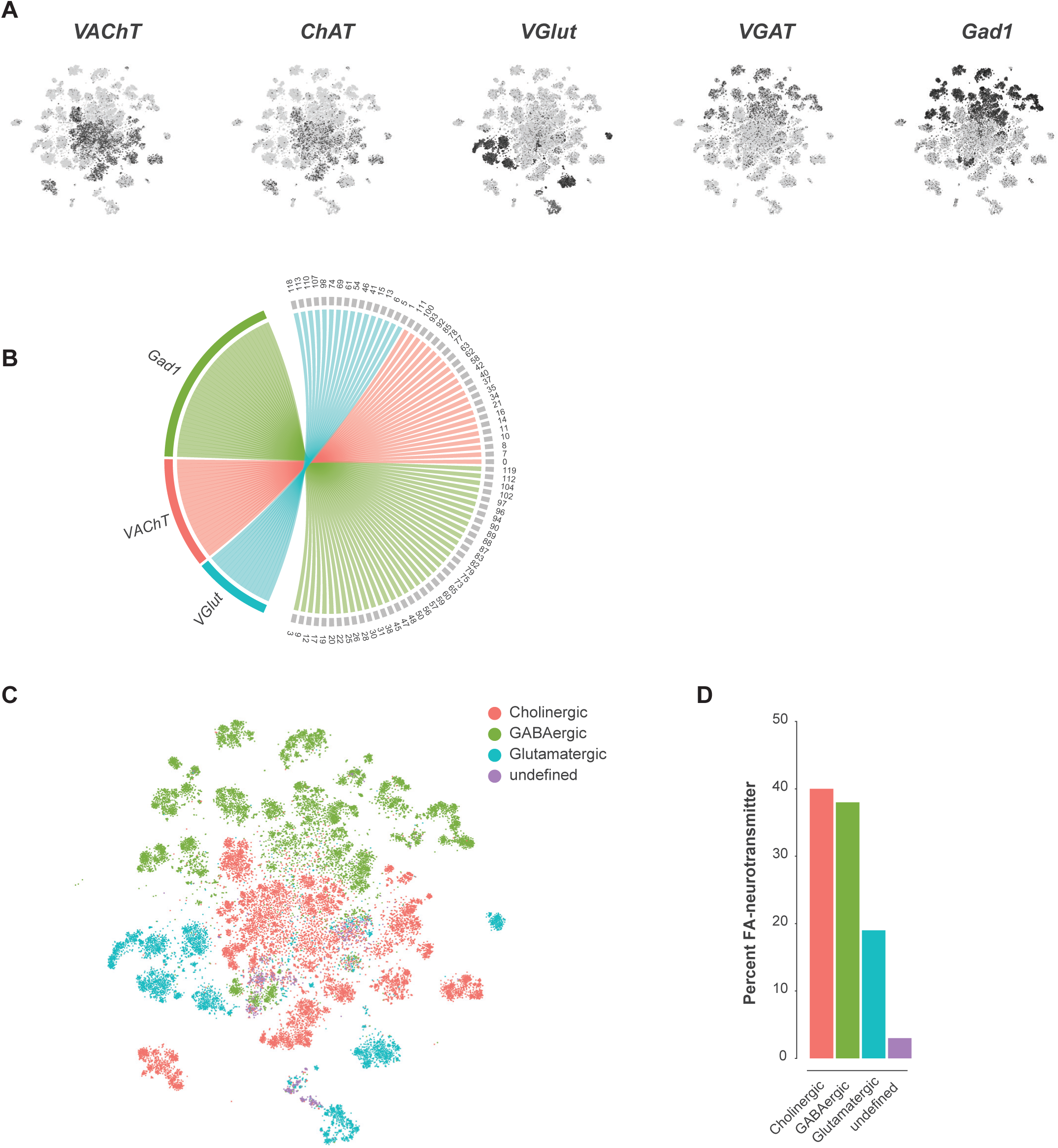
Fast-acting neurotransmitter usage in the VNC. A. t-SNE plots showing distribution of cells expressing biomarkers for the fact-acting neurotransmitters acetylcholine (*VAChT, ChAT*), glutamate (*VGlut*) and GABA (*VGAT, Gad1*). B. Chord diagram showing fast-acting neurotransmitter cluster markers (left, in color) with the clusters for which they are significantly enriched (right, in grey). C. t-SNE plot colored according to fast-acting neurotransmitter usage based on assigned cluster identity. D. Percentage of cells in the VNC assigned as releasing distinct fast-acting neurotransmitters.

### Monoaminergic neurons

Cells that produce and likely release monoamines in the VNC can be identified based on expression of the *Vesicular monoamine transporter* (*Vmat*) gene (Greer et al., 2005). To examine the anatomical distribution of *Vmat* expressing nuclei and projections in the VNC we combined *Vmat^GAL4^* with *UAS-Watermelon* (WM) (Figure 7A). *Vmat^GAL4^* labels 115 ± 6.3 cells distributed across every neuromere of the VNC (n = 9, data not shown). Each neuromere has a main cluster of cells located on the ventral surface, near the midline, consistent with known developmental origins of monoaminergic neurons. Serotonin- and dopamine-producing neurons are born from the ventromedially located paired neuroblast 7-3, and octopamine-producing neurons are born from the ventral midline unpaired median neuroblast (Bossing et al., 1996; Schmid et al., 1999). Projections of *Vmat-*labeled neurons formed thick fascicles of fibers projecting from each cluster to the dorsal surface of the VNC (Figure 7A, sagittal view). Each thoracic neuromere has one main fiber tract, whereas the ANm has 8 parallel tracts, one for each abdominal segment. Despite the relatively small number of *Vmat* expressing cells, they project throughout the VNC and densely innervate the entire neuropil. Additional *Vmat* nuclei associated with the VNC are consistent with its reported expression in perineural surface glia (DeSalvo et al., 2014). Surface glial *Vmat* expressing cells can be seen in cluster 98 of our primary t-SNE plot (Figure 1 and Figure supplement 1).

**Figure 7:**
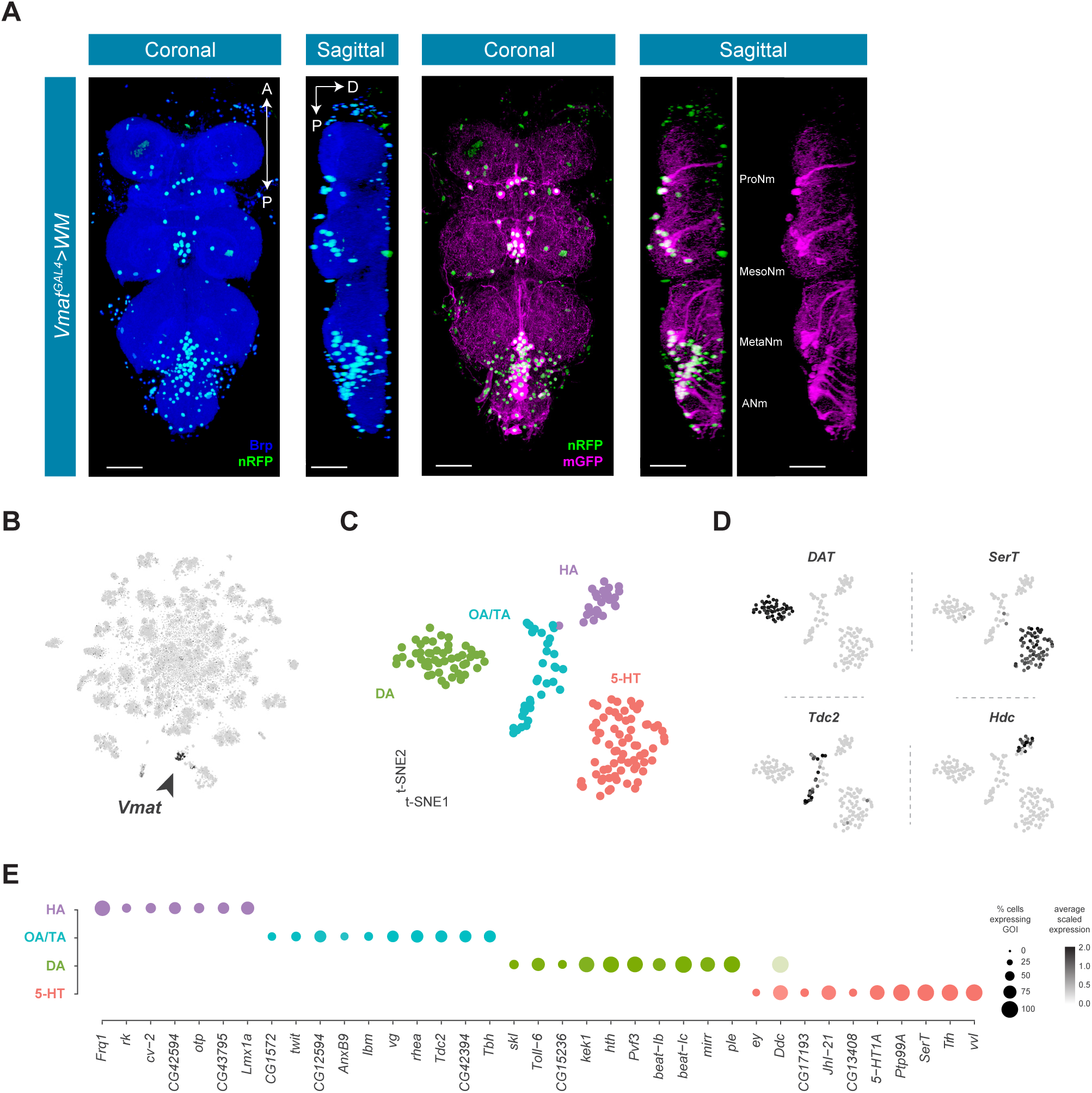
Identification and characterization of monoaminergic cell sub-types. A. Visualisation of *Vmat* expressing cells in the 5-day old adult VNC. Coronal and sagittal views of *Vmat^GAL4^* driving expression of *UAS-WM*, enabling the visualization of cell nuclei (nRFP; magenta) and neuronal morphology (mGFP; green). Neuropil is counterstained (Brp; blue). A, Anterior, P, posterior, D, dorsal; scale bars = 50 μm. B. t-SNE plot of *Vmat* expression in black, intensity is proportional to the scaled log-normalized expression level. *Vmat* enriched Cluster 72 is highlighted with arrowhead. C. t-SNE plot showing sub-clustering analysis of *Vmat^+^* cells from clusters 72 and 84. Four sub-clusters are identified representing dopaminergic (DA), octopaminergic/tyraminergic (OA/TA), histaminergic (HA), and serotonergic neurons (5-HT). D. Expression of established monoaminergic subtype-specific biomarkers used to determine cluster identity. *Histidine decarboxylase* (*Hdc*) labels Histamine (HA) neurons, *Tyrosine decarboxylase 2* (*Tdc-2*) labels Tyramine (TA) and Octopamine (OA) neurons, *Dopamine transporter* (*DAT*) labels Dopamine (DA) neurons and *Serotonin transporter* (*SerT*) labels Serotonin (5-HT) neurons. E. Dot plot of the top genes in each monoaminergic sub-type based on fold-enrichment (Figure 8-source data 1). Size of dots represent percent of cells in cluster expressing gene of interest (GOI); intensity of color reflects average scaled expression (0-2).

Neurons that express *Vmat* co-cluster in the single-cell data (Clusters 72 and 84, Figure Supplement 1 and Figure 7B). To define the specific monoaminergic identity of *Vmat* expressing neurons, we sub-clustered these *Vmat*^+^ cells from clusters 72 and 84 (Figure 7C) and identified distinct groups synthesizing specific monoamines, as defined by expression of known biosynthetic and transporter markers (Figure 7D) (Martin and Krantz, 2014).

### Histaminergic neurons

Histamine (HA) is well established as the primary fast neurotransmitter of adult photoreceptors in many insects, including *Drosophila* (Hardie, 1987; Nässel et al., 1988). The VNC contains 18 ventral HA-immunoreactive neurons with extensive axons and projections, six in the thoracic neuromeres and twelve in the abdominal neuromere (Buchner et al., 1993; Nässel et al., 1990). Potential new markers for HA neurons, distinguishing them from other monoaminergic neurons, include *Frequenin 1* (*Frq1*), a Ca^2+^-binding protein that regulates neurotransmitter release (Dason et al., 2009), and *CG43795*; an uncharacterized GPCR predicted to have Glutamate/GABA receptor activity (Agrawal et al., 2013).

### Tyraminergic and Octopaminergic neurons

Tyramine (TA) and Octopmamine (OA) neurons innervate and modulate many tissues throughout the fly, including female and male reproductive systems, skeletal muscles, and sensory organs (Pauls et al., 2018; Rezával et al., 2014). *Tdc2* encoded tyrosine decarboxylase catalyzes the synthesis of TA (Cole et al., 2005), which can also be converted to OA by *Tbh* encoded Tyramine beta-hydroxylase (Monastirioti et al., 1996). All *Tdc2* expressing neurons in the VNC also express *Tbh* suggesting that they are likely to be octopaminergic, consistent with a previous report using *Tdc2^GAL4^* in the VNC (Pauls et al., 2018). Prior work in the larval VNC suggested that the TA:OA ratio might be altered in a state-dependent manner (Schutzler et al., 2019).

### Dopaminergic neurons

Dopamine (DA) is a critical neuromodulator controlling learning and state-dependent plasticity in the fly brain (Cognigni et al., 2018). Dopamine and DA neurons in the VNC have been implicated in motor behaviors such as grooming and copulation (Crickmore and Vosshall, 2013; Yellman et al., 1997). Novel DA markers in the VNC include *beat-Ib* and *beat-Ic*, a likely recent duplication in the *beat* family of genes which act as heterophilic cell-cell adhesion molecules (Pipes et al., 2001). VNC DA neurons overexpress *PDGF- and VEGF-related growth factor* (*Pvf3)* and *kekkon 1* (*kek1*) which encodes a transmembrane protein that binds the EGF receptor and controls the activity of this pathway. Adult midbrain DA neurons also overexpress these genes (Croset et al., 2018), suggesting a possible common role for *Pvf3* and *kek1* in DA neuron function. The *Toll-6* neurotrophin-like receptor implicated in neuronal survival and motor-axon targeting (McIlroy et al., 2013), appears specifically enriched in VNC DA neurons.

### Serotonergic neurons

Serotonin (5-hydroxytryptamine, 5-HT) releasing neurons within the thoracic neuromeres of the VNC modulate walking speed in a context-independent manner as well as in response to startling stimuli (Howard et al., 2019). In the abdominal ganglion, two clusters of sexually dimorphic neurons expressing 5-HT and *fruitless* (approximately ten cells per cluster in males) innervate the internal reproductive organs. These abdominal clusters control transfer of sperm and seminal fluid during copulation (Billeter et al., 2006; Lee and Hall, 2001; Lee et al., 2001). 5-HT neurons in the VNC express the 5-HT receptor *5-HT1A*, suggesting auto-regulatory/autocrine control of 5-HT neurons. VNC 5-HT neurons also express the amino acid transporter encoded by *juvenile hormone inducible 21 (JhI-21)* which is required for 5-HT-dependent evaluation of dietary protein (Ro et al., 2016). Finding *JhI-21* expression in 5-HT neurons in the VNC, therefore suggests that some 5-HT neurons may directly sense circulating amino acids.

### A transcription factor code for monoaminergic neurons

The top group of genes defining each subclass of monoaminergic neurons types also included unique expression of Homeobox-containing TFs (Bürglin and Affolter, 2016) (Figure 7E). *orthopedia* (*otp*) and the *LIM homeobox transcription factor 1 alpha* (*Lmx1a*) define the HA cluster. DA neurons are enriched for the Hox co-factor *homothorax* (*hth*) as well as the Iroquois homeobox TF *mirror* (*mirr*). Lastly, 5-HT neurons express the homeobox TFs *ventral veins lacking* (*vvl*), *eyeless* (*ey*) and *Lim3*. Although these different combinations of homeobox genes likely act in these cells to specify position-specific patterning decisions and wiring specificity during development, it is also possible that they contribute to the function of mature monoaminergic neurons.

### Peptidergic neurons

Neuropeptides are the largest family of signaling molecules in the nervous system and act as important regulators of development, physiology, and behavior (Nässel and Zandawala, 2019). We analyzed neuropeptide expression in the adult VNC by looking at the expression of genes encoding neuropeptide precursors, from which active neuropeptides are derived. We identified 28 neuropeptides that are expressed in at least one cell, at a level of 10 or more transcripts per cell (Figure 8A). Some neuropeptides are known to be co-expressed with FANs, where they serve to increase signaling flexibility within neural networks (Croset et al., 2018; Nässel, 2018; Nusbaum et al., 2017). We found that in the VNC some neuropeptide genes are predominantly co-expressed with particular FANs, e.g. *sNPF* and *spab* in cholinergic neurons, *MIP* and *CCHa2* in GABAergic neurons, and *Ilp7* and *Proc* in glutamatergic neurons (Figure 8A; Figure 6). A few neuropeptides were also co-expressed in the same cells, e.g. *AstC* and *CCHa2* in GABAergic cluster 94. Some of these associations are similar to what was observed previously in midbrain single-cell data (Croset et al., 2018), e.g. *sNPF* and *spab* in cholinergic neurons, but others are strikingly different. *AstC* and *CCHa2* are robustly expressed in GABAergic neurons in the VNC but are notably absent from *Gad1* expressing neurons in the midbrain.

**Figure 8:**
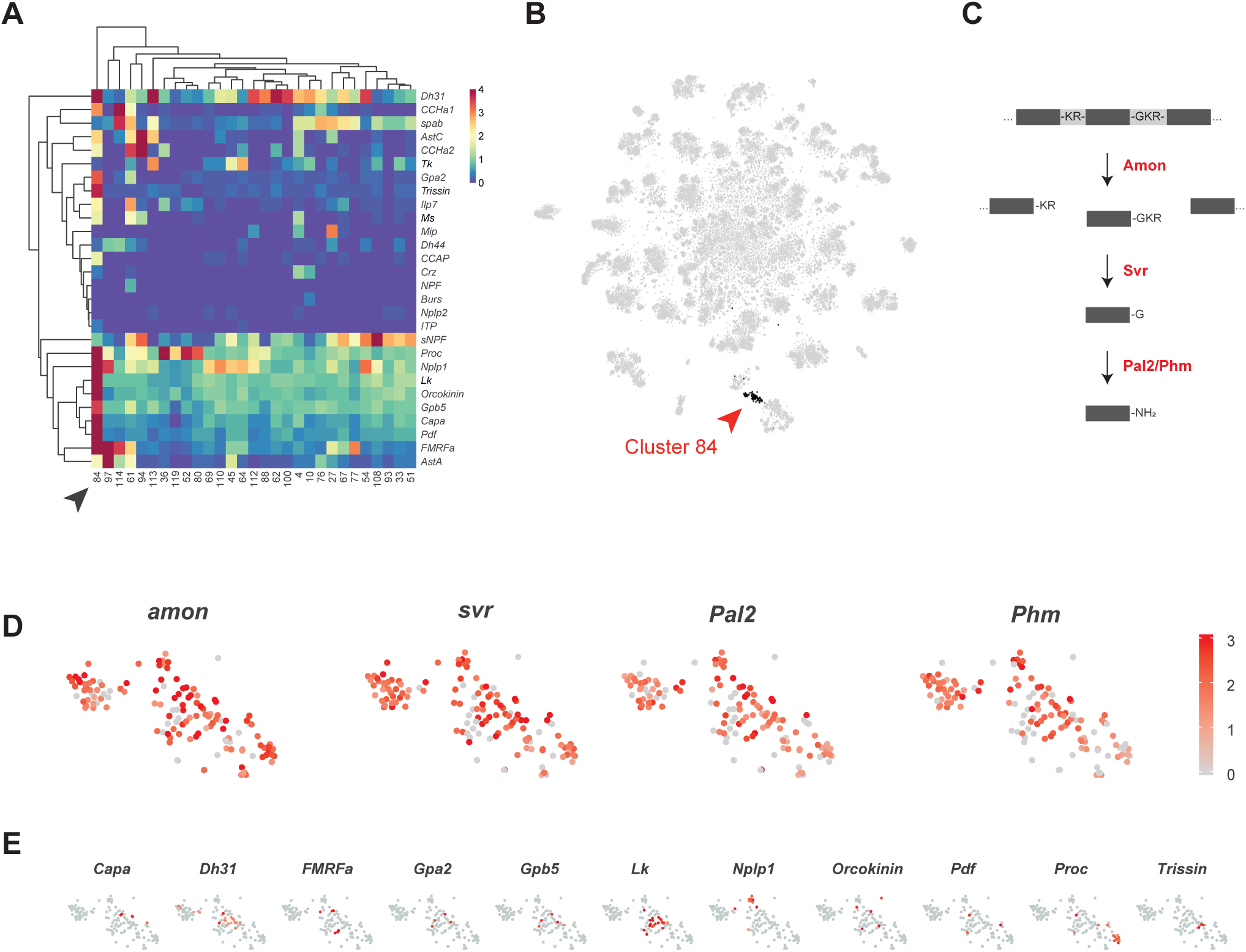
Identification of neuropeptide expressing cells. A. Heatmap of the mean log-normalized expression (0-4), by cluster (bottom), of neuropeptide genes (right). Only clusters with at least one neuropeptide gene expressing in excess of mean log-normalized value of 10 are shown. Only neuropeptide genes that were expressed in at least one cells having >10 transcripts per cell are shown. Arrowhead highlights cluster 84. Dendrograms represent hierarchical clustering. B. t-SNE plot highlighting neuropeptide expressing cluster 84. C. Schematic of neuropeptide processing steps highlighting enzymes (in red) identified as enriched markers for cluster 84. Propeptides are cleaved by the pro-hormone convertase *Amontillado* (*Amon*); the carboxypeptidase *Silver* (*Svr*) then removes the C-terminal cleavage sequence. C-terminal amidation occurs through the combined actions of *Peptidyl-α-hydroxyglycine-α-amidating lyase 2* (*Pal2*) and *Peptidylglycin-α-hydroxylating monooxygenase* (*Phm*) (reviewed in Pauls et al., 2014) D, E. Expression of neuropeptide processing enzymes (D) and multiple neuropeptide genes (E) in t-SNE cluster 84. Intensity of red is proportional to the log-normalized expression level.

Some neuropeptides are also expressed in ‘neurosecretory’ neurons that lack FANs. Peptides released from these cells into circulation provide neuroendocrine signals, whereas those released into the nervous system have neuromodulatory function. Cells in cluster 84 exhibit several characteristics of neurosecretory cells. The TF *dimmed* (*dimm*) is required for the differentiation of neurosecretory cells (Hamanaka et al., 2010; Hewes et al., 2003; Park et al., 2008). Although *dimm* is detected at very low levels and in few cells in our dataset, it shows a bias to cluster 84 (10.7% of cells express *dimm* vs. 0.03% elsewhere). Cluster 84 is also particularly enriched for neuropeptide expression (Figure 8A and B) and lacking strong expression of FAN or monoaminergic neuron markers (Figure 6C; Figure 7B). They also express several genes involved in the neuropeptide processing (see Figure 8C and D; Figure 1-source data 1). Most neuropeptides are expressed in non-overlapping subsets of the cluster (figure 8E), suggesting that a particular neuropeptide does not drive the unifying identity of the cluster, but rather that it is generically peptidergic. The *Glycoprotein hormone alpha 2* (*Gpa2*) and *Glycoprotein hormone beta 5* (*Gpb5*) genes, whose peptides form heterodimers, are co-expressed in the same cells in this peptidergic cluster (Sellami et al., 2011).

### High neuropeptide gene expression reveals important technical considerations

When investigating neuropeptide expressing cluster 84 (NP cluster) we noticed that the median number of transcripts in the cluster was 5277, which was far greater than that observed in the data set as a whole, 2497 (Figure 8-Figure supplement 1A). This large number of transcripts is due, in part, to high expression levels of the neuropeptide genes themselves. For example, the maximum expression level of *Leucokinin* (*Lk*) was 2629 transcripts in a single cell (Figure 8-Figure supplement 1B), whereas the maximum expression level of an average gene was 11 transcripts per cell. Amongst neuropeptide genes, *Lk* is not an anomaly, as 18 of the top 40 most highly expressed genes per cell encode neuropeptides (Figure 8-Figure supplement 1C). In some cases, they represent 40% of a cell’s entire transcriptional output (Figure 8-Figure supplement 1D).

For all previous data analyses, we used an upper limit of 10,000 total transcripts (nUMI) per cell to remove potential ‘doublets’, where two cells were co-encapsulated, and their transcriptional profiles merged. Given the high expression seen for neuropeptide genes, we repeated our analysis without this cut-off, revealing that many cells in the neuropeptide (NP) and *Proctolin+* motor neuron (*Proc^+^* MN) clusters express more than 10,000 transcripts (Figure 8-Figure supplement 2). It is worth considering the filtering cut-off when studying cells with high transcriptional output, and the genes expressed therein, whose expression can exceed the threshold. For example, the maximum observed expression of *Orcokinin* was 23,092 transcripts.

Another important consideration when investigating cells expressing genes at very high levels, as many neuropeptide cells do, is the consequence of these cells rupturing during the dissociation process. Once ruptured, large numbers of neuropeptide transcripts could be present in the ambient solution, leading to background levels of neuropeptide transcripts being ‘picked up’ with non-expressing cells. Our data for the neuropeptide *Orcokinin* illustrates this point (Figure 8-Figure supplement 3). Orcokinin is expressed in just 5 neurons in the adult VNC, two pairs of neurons in thoracic neuromeres, with additional expression in one unpaired neuron in the abdominal neuromere (Chen et al., 2015). In our data, 5 cells in the NP cluster show *Orcokinin* expression at a level of more than 100 transcripts (Figure 8-Figure supplement 3). However, almost 8000 cells, across all clusters, also appear to express *Orcokinin* at a level of just 1-10 transcripts (Figure 8-Figure supplement 3). We speculate that the low-level expression outside the NP cluster reflects background levels, due to rupture of *Orcokinin* expressing cells, while the expression above 100 transcripts within the NP cluster is *bona fide*.

### Non-neuronal cell types: Glia

Glia are key regulators of nervous system physiology maintaining the concentration of chemicals in the extracellular environment. We identified glia cells in our VNC data using the established glial markers *reversed polarity* (*repo*) (Xiong et al., 1994) and the long non-coding RNA *MRE16* (Davie et al., 2018) (Figure 9-Figure supplement 1). Three clusters, representing 3.6% of the total cells, were highly enriched for *repo* and *MRE16.* In addition, these clusters lacked expression of neuronal markers, such as *elav*, *nSyb*, and *noe* (Figure 9-Figure supplement 1). Two additional clusters contained cells that were positive for both glial markers and cells positive for neuronal markers, suggesting mixed populations (Figure 9-Figure supplement 1). To define specific glial cell types, we performed a sub-clustering analysis of the five glial clusters (Figure 9A), while removing any cell with a neuronal signature. This sub-clustering revealed four distinct glial subpopulations that could be classified based on enriched expression of established markers (Figure 9B): *astrocytic leucine-rich repeat molecule* (*alrm*) for astrocytes (Doherty et al., 2009), *I’m not dead yet* (*Indy*) for surface glia (Knauf et al., 2002), *wrapper* for cortex glia (Noordermeer et al., 1998), and *Excitatory amino acid transporter 2* (*Eaat2*) for ensheathing glia (Stahl et al., 2018).

**Figure 9:**
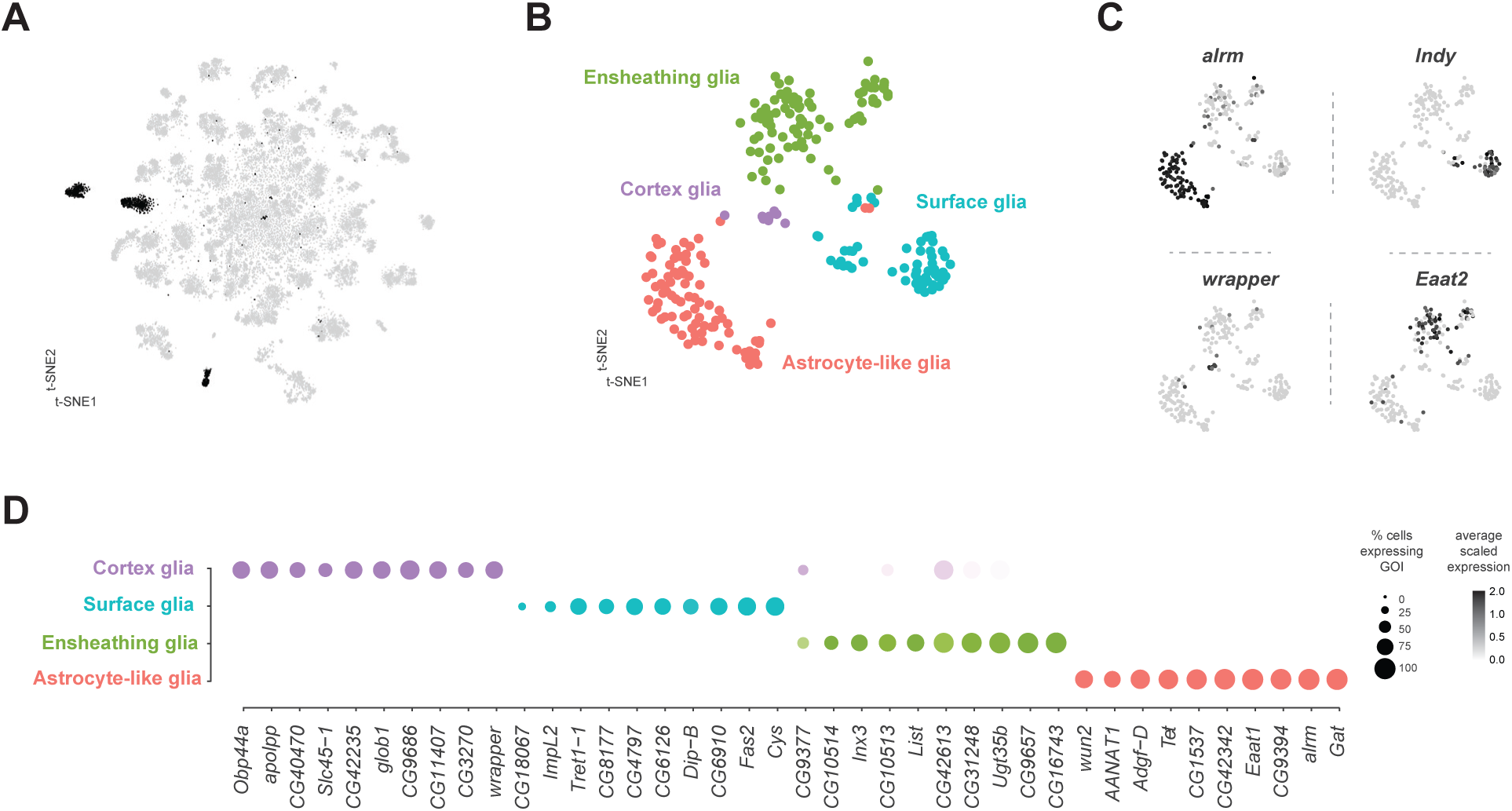
Identification and characterization of glial cell sub-types. A. t-SNE plot highlighting glial cell clusters. Clusters 23, 24, 70, 98, and 106 are shown in black, as they were determined to be glia based on differential expression of established glial and neuronal biomarkers (Figure 10-Figure supplement 1). B. t-SNE plot showing sub-clustering analysis of glial cells. Distinct clusters are color-coded and named. C. Expression of established glial subtype-specific biomarkers used to determine cluster identity. *astrocytic leucine-rich repeat molecule* (*alrm*), for astrocytes; *I’m not dead yet* (*Indy*) for surface glia; *wrapper* for cortex glia; *Excitatory amino acid transporter 2* (*Eaat2*) for ensheathing glia. D. Dot plot of the top 10 genes in each glial sub-type based on fold-enrichment (Figure 10-source data 1). Size of dots represent percent of cells in cluster expressing gene of interest (GOI); intensity of color reflects average scaled expression (0-2).

### Surface glia

Consistent with the established metabolic role of glia in the brain, many solute carrier (SLC) membrane transporters are enriched in glial cell types (Figure 9D; Figure 9-source data 1). Surface glia, which form the blood-brain barrier (BBB), metabolically insulate the nervous system from the hemolymph. The BBB also subserves the considerable energetic demands of the nervous system (Laughlin et al., 1998), by efficiently transporting sugars, ions, and metabolites between the hemolymph and brain (Limmer et al., 2014). Surface glia strongly express the previously described surface glia marker SLC2 *Trehalose transporter 1-1* (*Tret1-1*), which as the name suggests transports trehalose, the main carbohydrate in the hemolymph, into the nervous system (Volkenhoff et al., 2015). Surface glia also contain high levels of a putative SLC2 sugar transporter *CG4797*, previously reported to be expressed in the perineural glia of the optic lobe (Figure 9D) (Konstantinides et al., 2018). Both of these sugar transporters are proton-dependent and therefore, efficient sugar transport requires glial cells to have lower H^+^ concentration than the hemolymph (Kikuta et al., 2012). We found enriched expression of multiple V-type ATPase H^+^ pumping genes (p-value = 3.9×10^-9^ based on DAVID analysis; Figure 9-source data 1) (Chintapalli et al., 2013) in surface glia. Expression of the SLC bicarbonate transporter *CG8177*, which has been shown to reduce extracellular pH (Overend et al., 2016), and the *Ecdysone-inducible gene L2* (*ImpL2*) were also enriched in surface glia (Figure 9D). *ImpL2* antagonizes *insulin-like peptide 2* (*Dilp2*) and inhibits insulin/insulin-like growth factor signaling (Honegger et al., 2008) suggesting that the BBB is responsive to the metabolic demands of the fly.

### Astrocytes and Ensheathing glia

Over 70% of the VNC astrocyte cell-specific markers are shared with those in astrocytes in the midbrain (Croset et al., 2018). This similarity suggests that there is minimal regional specialization of astrocyte identity and function within the CNS. All astrocytes and ensheathing glia in the thoracic VNC are derived from the same lineages as leg motor neurons (Enriquez et al., 2018). However, we did not find evidence of a lineage-specific transcriptional code in these glial cell types (Figure 9-source data 1). Astrocytes instead, strongly express the paired-like homeobox TF *CG34367*, previously reported to be expressed in astrocytes throughout development (Huang et al., 2015). In contrast, ensheathing glia express the POU homeobox TF *ventral veins lacking* (*vvl*) (Anderson et al., 1995). These TFs can, therefore, be considered as novel markers for these adult VNC cell types. The rarity of TF markers is in accord with the finding that the morphological specificity of motor neurons depends on a unique TF code, whereas astrocytes and ensheathing glia show plasticity in their morphologies dependent on their position, rather than a distinct TF code (Enriquez et al., 2018).

### Non-neural ‘contamination’

We identified two clusters of cells that represent contamination during the dissection process (Figure 9-Figure supplement 1B). Cluster 99 was determined to be salivary gland tissue, since the cluster markers for cluster 99 contains 8 of the top 10 genes enriched in salivary gland tissue in the FlyAtlas 2 dataset (Figure 1-source data 1) (Leader et al., 2018). Cluster 116 appears to be sperm as many of the markers for this cluster are unique to the testis and sperm (Figure 1-source data 1) (Witt et al., 2019).

## Discussion

As scientists try to categorize nervous systems at higher resolution, the question arises what precisely constitutes a cell type? The goal itself seems straightforward in principle, to find a way to define different groups of cells that carry out distinct tasks. Single-cell mRNA sequencing techniques provide a possible route to answering this question by allowing the neuronal transcriptome of thousands of individual cells within a complex nervous system to be collected in parallel. To this end we generated a single-cell transcriptional atlas that reveals extensive cellular diversity in the adult *Drosophila* ventral nerve cord. In combination with previous single-cell data from the antennal lobe, optic lobe, and brain (Croset et al., 2018) (Davie et al., 2018; Konstantinides et al., 2018; Li et al., 2017), our data contribute towards a comprehensive cell atlas representative of the entire adult central nervous system.

Distinguishing between some cell types such as neurons and glia is relatively straightforward, but the extent of neuronal diversity provides a real challenge. Different neuronal types transmit particular neurotransmitters, neuropeptides and monoamines and they also respond to a variety of these signals using their complement of cell-surface receptors. Certain neurons might also express unique ion-channels and cell-signaling cascades that provide the cell with a range of electrophysiological characteristics, and potential mechanisms of plasticity. In principle we can also define neuronal cell types by characterizing their neuroanatomy - where the neurons are located and to which neurons they are pre- and post-synaptically connected. Since neurons acquire their anatomy through developmental programs it was not known whether this information would remain accessible in any form in the snapshot of the transcriptome of mature fully differentiated adult VNC. However, one of the clearest cluster-defining features of our VNC data is the abundance of transcription factors and cell-adhesion molecules that are classically thought of as being developmental.

The cluster-defining TFs are particularly useful for annotating the cell types in the VNC. Decades of work has investigated the development and structural organization of the VNC and has described roles for these genes in developmental specification of hemilineages (Baek and Mann, 2009; Birkholz et al., 2015; Bossing et al., 1996; Lacin et al., 2019; Lacin and Truman, 2016; Prokop and Technau, 1991; Schmid et al., 1999; Shepherd et al., 2016; Shepherd et al., 2019; Truman and Bate, 1988; Truman et al., 2004). Hemilineages are the functional units of the VNC and represent key organisational principals for connectivity. Overwhelming, the neuronal diversity observed in the VNC could be attributed to cells with shared developmental hemilineage histories. In principle defining the hemilineage identity of a cluster permits prediction of the location of a neuron, the neuropils its neurites innervate, and its likely motor outputs (Birkholz et al., 2015; Harris et al., 2015; Lacin et al., 2019; Shepherd et al., 2019; Venkatasubramanian and Mann, 2019). Cellular diversity within hemilineages was also apparent in our data. For example, we documented that *kn* and *twz* are expressed in distinct subsets of hemilineage 23B. 23B neurons project ventrally and innervate leg associated neuropils (Lacin and Truman, 2016; Shepherd et al., 2019) and their thermogenetic activation, in a headless fly, induces uncoordinated leg movement and a change in body posture (Harris et al., 2015). Generation of tools to functionally and anatomically investigate these subsets of hemilineage 23B based on the identification of *kn* and *twz* as novel sub-cluster markers, in addition to investigating the transcriptional diversity of these cells will facilitate a functional dissection of their specific roles in leg coordination and/or body posture.

We also found continued expression in hemilineages of neurodevelopment genes, such as *br*, *Imp*, and *pros*, which may signify cell birth order and therefore further distinguish cell types. Expression levels of *pros* indicate the birth order of post-embryonic leg motor neurons belonging to hemilineage 15B, that precedes muscle-specific innervation (Baek et al., 2013; Baek and Mann, 2009). Relative levels of *Imp* and *pros* in our predicted hemilineage 15B, can therefore be used to infer muscle-specific innervation along the leg. *Imp* and *pros* expression may also distinguish between embryonic born and post-embryonic born neurons. Mature (embryonic) neurons of the larval VNC show elevated *Imp* expression, whereas immature (post-embryonic) neurons have higher levels of *pros*, relative to each other (Etheredge, 2017). Dopaminergic neurons in the PAM1 cluster in the larva are the only dopaminergic neurons to be born post-embryonically (Hartenstein et al., 2017). Embryonic vs. post-embryonic identity can therefore also sometimes be used to identify neurons within a given class.

Fast-acting neurotransmitter identity is acquired in a hemilineage dependent manner, amongst post-embryonic neurons (Lacin et al., 2019), but the underlying transcriptional programs appear to be complex (Estacio-Gómez et al., 2019). We did not identify single TFs that globally defined FAN identity. Different neurons have been reported to use different TFs to define the same FAN identity in the adult fly optic lobes (Konstantinides et al., 2018) and in *C. elegans* (Hobert and Kratsios, 2019). In addition, a recent gene-profiling study of cholinergic, glutamatergic, and GABAergic neurons across development found that many FAN-specific TFs are transiently expressed at particular times, and only a few remain constant (Estacio-Gómez et al., 2019). As noted previously (Lacin et al., 2019), we found some TFs that are restricted to individual FAN-specific clusters. Some GABAergic clusters express *Dbx*, *vg*, *D*; some cholinergic neurons express *unc-4* and some glutamatergic neurons express *ems*. We also found *Lim3* to be expressed in cholinergic and glutamatergic neurons, but not in GABAergic neurons (Lacin et al., 2019). In contrast *Lim3* specifically regulates GABAergic cellular identity in the optic lobes (Konstantinides et al., 2018). Yet, consistent with the optic lobes, *Lim3* is only expressed in Notch^off^ hemilineages in the VNC (Li et al., 2017). Therefore, the TF code specifying FAN identity may change across development and also differ depending on the regional context.

We could place the cells from our data into the anatomical landscape of the VNC using their *Hox* gene expression. *Antp*, *Ubx*, *abd-A*, and *Abd-B*, which developmentally define the neuromeres of the VNC, continue to be expressed in the adult. These genes permit assignment of neuron profiles to the prothoracic, mesothoracic, metathoracic and abdominal neuromeres. Finding that established neuromere- and hemilineage-specific developmental markers are expressed in the 5-day old adult, suggests they may continue to act in these differentiated cells to maintain the distinct transcriptional profile underlying neuronal identity and function (Deneris and Hobert, 2014). It will therefore be interesting to investigate functional consequences of disrupting these potential maintenance programs.

Not all cells group by their developmental lineage, for example, monoaminergic cells formed a distinct cluster, despite originating from multiple different neuroblasts (Schmid et al., 1999). We also identified a cluster of neurosecretory cells that includes cells with different developmental origins (Park et al., 2008). High-level expression of neuropeptides in this cluster is consistent with features of neurosecretory biology. Neuropeptides are often produced by small numbers of cells, yet they can be broadly released and by volume transmission and/or secretion into the hemolymph act at a distance to modulate disparate neural circuits, physiological functions and behaviors (Nässel, 2018). The extraordinarily high-level expression of pre-pro-neuropeptide genes in these cells suggests that producing these molecules presents a considerable burden. The TF *dimmed* (*dimm*), which marks neurosecretory cells of this type, has been shown to promote neurosecretory identity and suppress FAN identity (Hewes et al., 2003; Park et al., 2008).

In this study, we have demonstrated the predictive power of the single-cell atlas of the VNC. Combining cell-specific expression with developmental information permitted the assignment of many clusters and cells to known neurons and anatomical locations in the VNC. The atlas also provides hundreds of new marker genes for each of these clusters and cell types. We assume that many of these novel genes will be important for the function of the relevant cells and will generate numerous new and testable hypotheses.

## Materials and Methods

### Resources Table

#### Primary Antibodies

Primary Antibodies:
| Antibody | Source | Identifier | Host species | Concentration |
| --- | --- | --- | --- | --- |
| AbdA (C-11) | Santa Cruz Biotechnology | sc-390990 | Mouse | 1:100 |
| AbdB (1A2E9)-s | DSHB | n/a | Mouse | 1:10 |
| Acj6-s | DSHB | n/a | Mouse | 1:20 |
| AntP 8C11-s | DSHB | n/a | Mouse | 1:20 |
| CadN (DN-Ex #8) | DSHB | n/a | Rat | 1:30 |
| nc82 (BRP) | DSHB | n/a | Mouse | 1:50 |
| GFP | Abcam | ab92456 | Chicken | 1:1200 |
| Ubx FP3.38-s | DSHB | n/a | Mouse | 1:20 |

#### Secondary Antibodies

Secondary Antibodies:
| Host species | Target Species | Conjugated fluorophore | Source | Identifier | Concentration |
| --- | --- | --- | --- | --- | --- |
| Goat | Mouse | AF 633 | Thermofisher Scientific | A21052 | 1:300 |
| Goat | Rat | AF 546 | Thermofisher Scientific | A11081 | 1:300 |
| Goat | Chicken | AF 488 | Thermofisher Scientific | A11039 | 1:300 |

#### Fly Strains and husbandry

The fly strain used for VNC analysis (+/w*; *UAS-Stinger*, *13XLexAop2-IVS-tdTomato.nls*/+; *dsx*^GAL4^/+) was a genetic cross between *w\**; *UAS-Stinger*, *13XLexAop2-IVS-tdTomato.nls* males and *+*; +; *dsx*^GAL4^ virgin females (Rideout et al., 2010). All flies were reared at 25°C in a 12:12 hr light:dark cycle on standard food at 40-50% relative humidity. Virgin males and females were collected and stored individually. Flies were aged 5 days post-eclosion at 25°C prior to dissection. Additional fly strains used in this study include:

wild-type *Canton-S*

*y^1^w*; Mi{Trojan-GAL4.0}Vmat[MI07680-TG4.0]* (BDSC:66806)

*y^1^w*; Mi{Trojan-GAL4.1Lim3}[MI03817-TG4.1]/SM6a* (BDSC:67450)

*y^1^w*; {Mi{Trojan-GAL4.2}kn[MI15480-TG4.2]/SM6a* (BDSC:67516)

*y^1^w*; Mi{Trojan-GAL4.2}twz[MI14153-TG4.2]/SM6a* (BDSC:76758)

*y^1^w* Mi{Trojan-GAL4.0}acj6[MI07818-TG4.0]/FM7c* (BDSC:77788)

*w*; {UAS-Stinger}* denoted as *UAS-nGFP* (Barolo et al., 2000)

*w*; UAS-myr-GFP-V5-P2A-H2B-mCherry-HA/TM3, Ser* (aka UAS-WM; Chang et al., 2019)

#### VNC single-cell sample preparation

The VNC dissociation protocol was carried out as described previously (Croset et al., 2018). 40 Male and 40 female VNCs (20 per sexed replicate) were individually dissected in toxin-supplemented ice-cold calcium- and magnesium-free DPBS (Gibco, 14190–086 + 50 µM D(−)-2-Amino-5-phosphonovaleric acid, 20 µM 6,7-dinitroquinoxaline-2,3-dione and 0.1 µM tetrodotoxin). Each replicate was then washed in 1 mL ice-cold toxin-supplemented Schneider’s medium (tSM: Gibco, 21720–001 + toxins, as above). VNCs were then incubated for 30 minutes in 0.5 mL of tSM containing 1 mg/mL papain (Sigma, P4762) and 1 mg/mL collagenase I (Sigma, C2674). VNCs were washed once more with tSM and subsequently triturated with flame-rounded 200 µL pipette tips. Dissociated VNCs were resuspended into 1 mL PBS + 0.01% BSA and filtered through a 10 µm CellTrix strainer (Sysmex, 04-0042-2314).

#### Data processing

Libraries were made using the Chromium Single Cell 3’ v2 kit from 10x Genomics. Cells were loaded in accordance with 10x Genomics documentation, with the aim of 5000-8000 cells per sample. The samples were sequenced with 8 lanes of Illumina HiSeq4000 by Oxford Genomics Centre. We obtained a mean of 20,550 reads per cell and mean sequence saturation of 71%. The fastq data were processed with Drop-seq_tools v2.1.0 (Macosko et al., 2015) and aligned to the *D. melanogaster* genome (R6.13) to generate digital expression matrices for each sample.

#### Data analysis with Seurat

The digital expression matrices were analyzed with the Seurat 2.3.4 R package (Satija et al., 2015). Genes expressed in fewer than 3 cells were removed. Cells with fewer than 1,200 UMI or more than 10,000 UMI were removed (unless otherwise mentioned). Cells with fewer than 200 genes, or more than 15% mitochondrial derived UMI were removed. Sex-specific replicates were merged, normalized, and scaled while regressing out variation due to replicate, nUMI, and proportion mitochondrial transcript. Low expression level, and low dispersion cut-offs of 0.001 were used to identify variable expressed genes in each sex. The intersection of these genes was used to perform a canonical correlation analysis (CCA) with the first 45 dimensions. t-distributed stochastic neighbor embedding (t-SNE) was performed, with perplexity of 30, theta of 0.05, and 20,000 iterations, on the data to reduce the dimensionality to 2 for visualization. Clusters were defined by a shared nearest neighbor method using 45 dimensions and a resolution of 12 (unless otherwise mentioned). Comparison of different cluster resolutions was evaluated with the ‘clustree’ package (Zappia and Oshlack, 2018). Positively enriched cluster markers were identified using a negative binomial distribution test with fold enrichment of at least 0.5 and a Bonferroni adjusted p-value of less than 0.05. Functionally related gene enrichment analysis was performed using DAVID (Huang et al., 2009). FlyAtlas 2 bulk RNA-seq data of the adult VNC was obtained from http://flyatlas.gla.ac.uk/FlyAtlas2/index.html (Leader et al., 2018). The convert IDs tool from Flybase was used to convert gene symbols and Flybase ID between release versions (http://flybase.org/convert/id).

#### Assigning fast-acting neurotransmitter identity

The following genes were used to assign fast-acting neurotransmitter (FAN) identity; *Vesicular acetylcholine transporter* (*VAChT*) and *Choline acetyltransferase* (*ChAT*) for acetylcholine, *Vesicular glutamate transporter* (*VGlut*) for glutamate, and *Vesicular GABA Transporter* (*VGAT*) and *Glutamic acid decarboxylase 1* (*Gad1*) for GABA. Cells from clusters with average scaled expression above 0.5 for *Gad1* or *VGAT*, and average scaled expression less than 0 for the other markers were all assigned to be GABAergic. Glutamatergic and cholinergic cells were assigned similarly. For clusters with an average expression less than 0.5 for all markers, cells were assigned FAN identity at a cell-by-cell level. We established different criteria for assigning cholinergic vs. glutamatergic or GABAergic since the cholinergic markers were expressed at much lower levels. If both *VAChT* and *ChAT* had a log-normalized expression greater than 0, then the cells was assigned to be cholinergic. Otherwise, if either *VAChT* or *ChAT* had a log-normalized expression greater than 2, then the cells was assigned to be cholinergic. Otherwise, if *VGlut* had a log-normalized expression greater than 2 and greater than either *VAChT* or *Gad1*, then the cells were assigned to be glutamatergic. Otherwise, if *Gad1* had a log-normalized expression greater than 2 and greater than either *VAChT* or *VGlut*, then the cells were assigned to be GABAergic. All remaining cells were labelled undefined.

#### Sub-clustering of monoaminergic cells

*Vmat* expressing cells from clusters 72 and 84 were sub-clustered. Identification of variable genes and canonical correlation analysis were performed as above. t-SNE was performed using the first 7 dimensions and a cluster resolution of 1.2 was used to identify clusters. Positively enriched cluster markers were identified as above.

#### Sub-clustering of glial cells

Clusters 23, 24, 70, 98, and 106 were identified as being predominantly glial cells. Cells from these clusters were sub-clustered. Any cell expressing *elav*, *nSyb*, *VAChT*, *VGlut*, or *Gad1* was removed. Identification of variable genes and canonical correlation analysis were performed as above. t-SNE was performed using the first 6 dimensions and a cluster resolution of 0.9 was used to identify clusters. Positively enriched cluster markers were identified as above.

#### R packages and plotting

Expression heatmaps were drawn using the ‘pheatmap’ package, correlation heatmaps were drawn using the ‘ggcorplot’ package, chord diagrams were drawn using the ‘circlize’ package (Gu et al., 2014), scatter plots and bar charts were drawn using ggplot2 in R. All plots were edited in Adobe Illustrator.

#### Phylogeny

The multiple sequence alignment (not shown) and the phylogenetic tree were created with Clustal Omega (Sievers and Higgins, 2018) and FigTreeV1.4.4 (http://tree.bio.ed.ac.uk/software/figtree/), respectively.

#### Immunohistochemistry

Flies were reared at 25°C and aged for 5 days prior to dissection and staining as per Rideout et al., (2010) with the following modifications: VNC samples were fixed in 4% PFA (40 mins at room temperature) immediately following dissection, to maintain tissue integrity and minimize cell loss. Samples were pre-incubated in 5% NGS overnight at 4°C. Samples were then incubated with primary antibodies for 3 days at 4°C, followed by an overnight incubation in secondary antibodies at 4°C. Primary antibodies used were: mouse mAb nC82 (1:50), mouse anti-AbdA (1:100, C-11), mouse anti-AbdB (1:10, 1A2E9-s), mouse anti-Acj6-s (1:20), mouse anti-AntP (1:20, 8C11-s), mouse anti-Ubx (1:20, FP3.38-s), rat anti-CadN (1:30, DN-Ex #8;) from DSHB, Univ. of Iowa. Additional primary antibodies used were chicken anti-GFP (1:1200, Abcam) and anti-AbdA (1:100, C-11 Santa Cruz Biotechnology). Secondary antibodies used included: anti-chicken Alexa Fluor488, anti-mouse Alexa Fluor633, anti-rat Alexa Fluor546, (1:300, Invitrogen Molecular Probes, Carlsbad, CA). Samples were left in 70% Glycerol/ 30% PBT overnight at 4°C prior to mounting with Vectashield (Vector Labs) and imaged with a Leica SP5 Microscope. Stacks of optical sections were generated at 0.5 µm intervals. Images were processed in Imaris 8.2.1 (Bitplane Scientific, AG, Zürich).

## Acknowledgements

We thank Tetsuya Nojima and Annika Rings for dissection assistance. We thank David Shepherd, Deniz Erezyilmaz, and members of the Goodwin lab for helpful discussions and critical reading of the manuscript. We thank the Bloomington Stock Center for flies. We thank Josh Dubnau for the UAS-WM line. Sequencing was carried out by The Oxford Genomics Centre. We thank Deborah J. Andrew and Caitlin D. Hanlon for assistance generating a Figure of the GPCRs. S.W. is supported by a Wellcome Principal Research Fellowship (200846/Z/16/Z) and ERC Advanced Grant (789274). This work was funded by a Wellcome Investigator Award (106189/Z/14/Z) to S.F.G, and a Wellcome Collaborative Award (209235/Z/17/Z) to S.F.G and S.W.

## Author Contributions

Aaron M. Allen, Conceptualization, Data curation, Formal analysis, Validation, Investigation, Visualization, Methodology, Writing—original draft, Writing—review and editing; Megan C. Neville, Conceptualization, Formal analysis, Validation, Investigation, Visualization, Methodology, Writing—original draft, Writing—review and editing; Sebastian Birtles, Formal analysis, Validation, Investigation, Visualization, Methodology, Writing— original draft, Writing—review and editing; Vincent Croset, Methodology, Writing—review and editing; Christoph D. Treiber, Methodology, Formal analysis; Scott Waddell, Conceptualization, Supervision, Funding acquisition, Writing—review and editing; Stephen F. Goodwin, Conceptualization, Supervision, Funding acquisition, Writing—original draft, Writing—review and editing

---

**Figure 1—source data 1: List of marker genes for the 120 clusters shown in Figure 1B.**

Table showing the average log-fold change values of cluster-discriminative marker genes, including adjusted p-values. pct.1 is the proportion of cells that express the gene in the cluster, pct.2 is the proportion of cells that express the gene in the other clusters. Predicted neurotransmitter usage and cell types represented in each cluster are specified where appropriate.

**Figure 1-Figure supplement 1:**
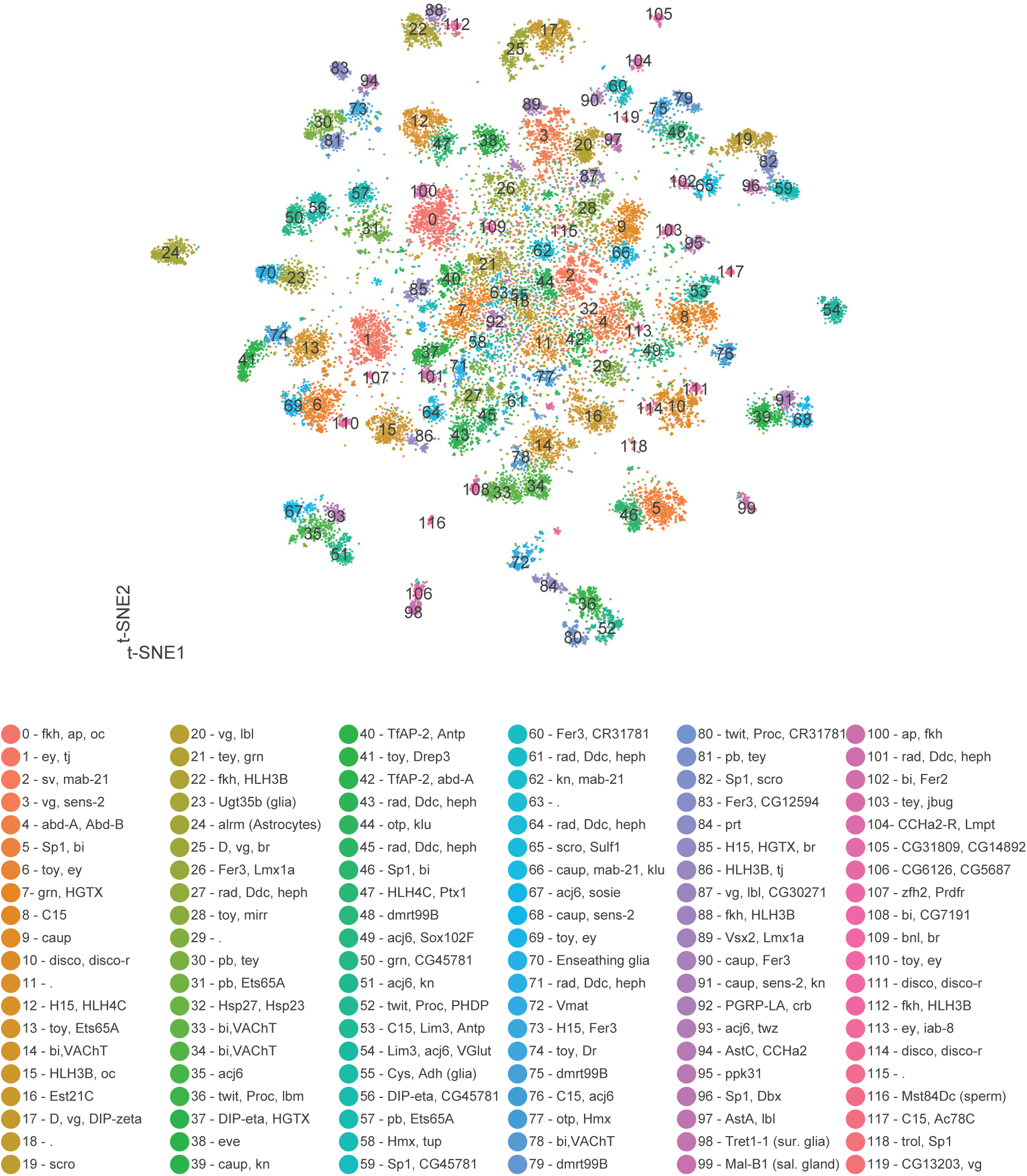
t-SNE plot of 5-day old adult VNC. t-SNE of VNC cells; each cluster is defined by a unique number and color. Where appropriate, combinations of genes uniquely identify each cluster are shown.

**Figure 1-Figure supplement 2:**
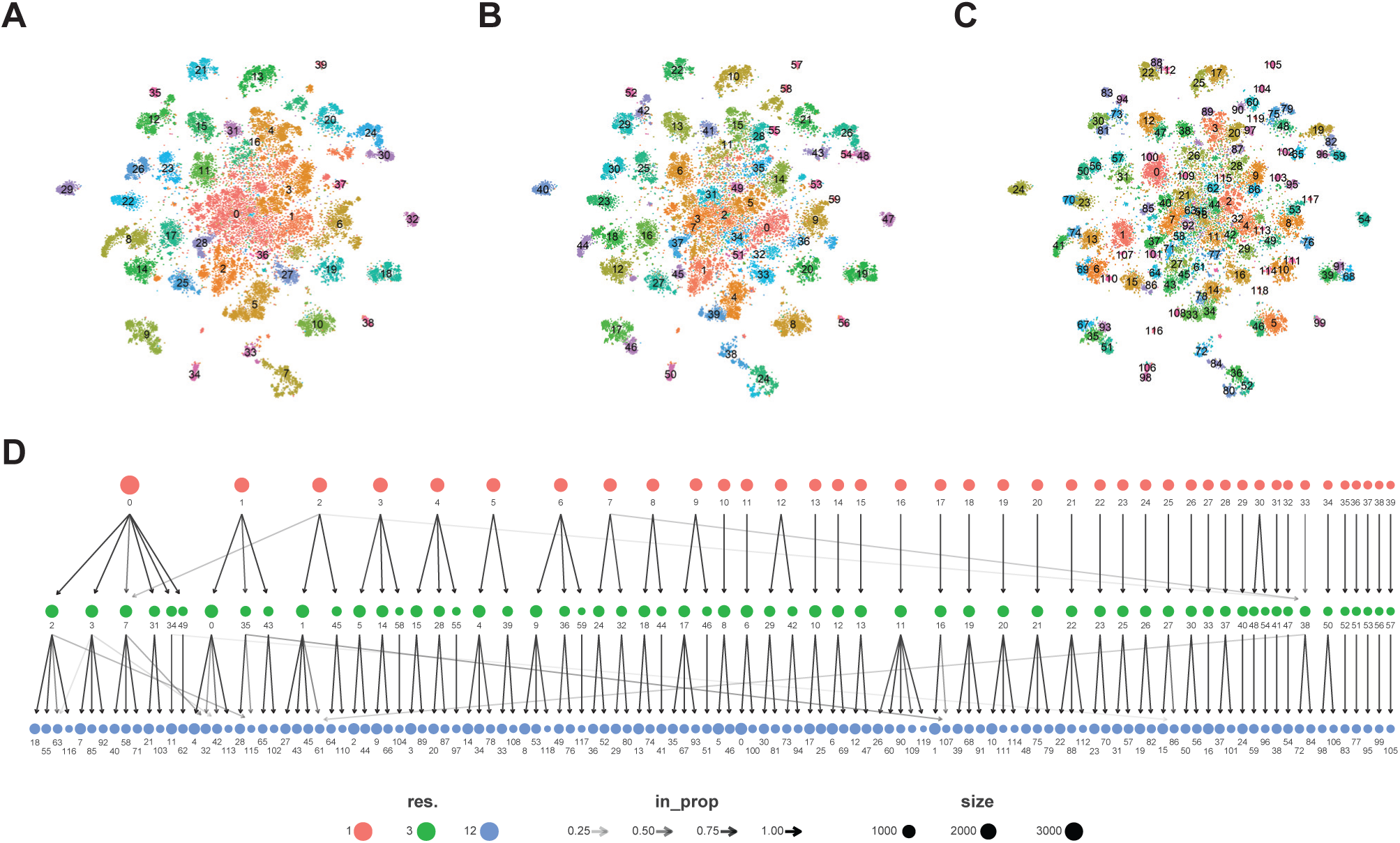
Cluster resolution analysis of adult VNC dataset. A-C. t-SNE plots with a cluster resolution of 1.0 (A), 3.0 (B), or 12.0 (C); Individual clusters are defined by unique numbers and colors. D. Dendrogram of the relationship between cluster (shown as numbers) at different cluster resolutions (color circles). The shade of the arrows connecting clusters between resolutions represent the proportion of the cluster at the higher resolution (arrowhead) found in the cluster from the lower resolution (arrow start). The number of cells in each cluster is represented by their relative size.

**Figure 1-Figure supplement 3:**
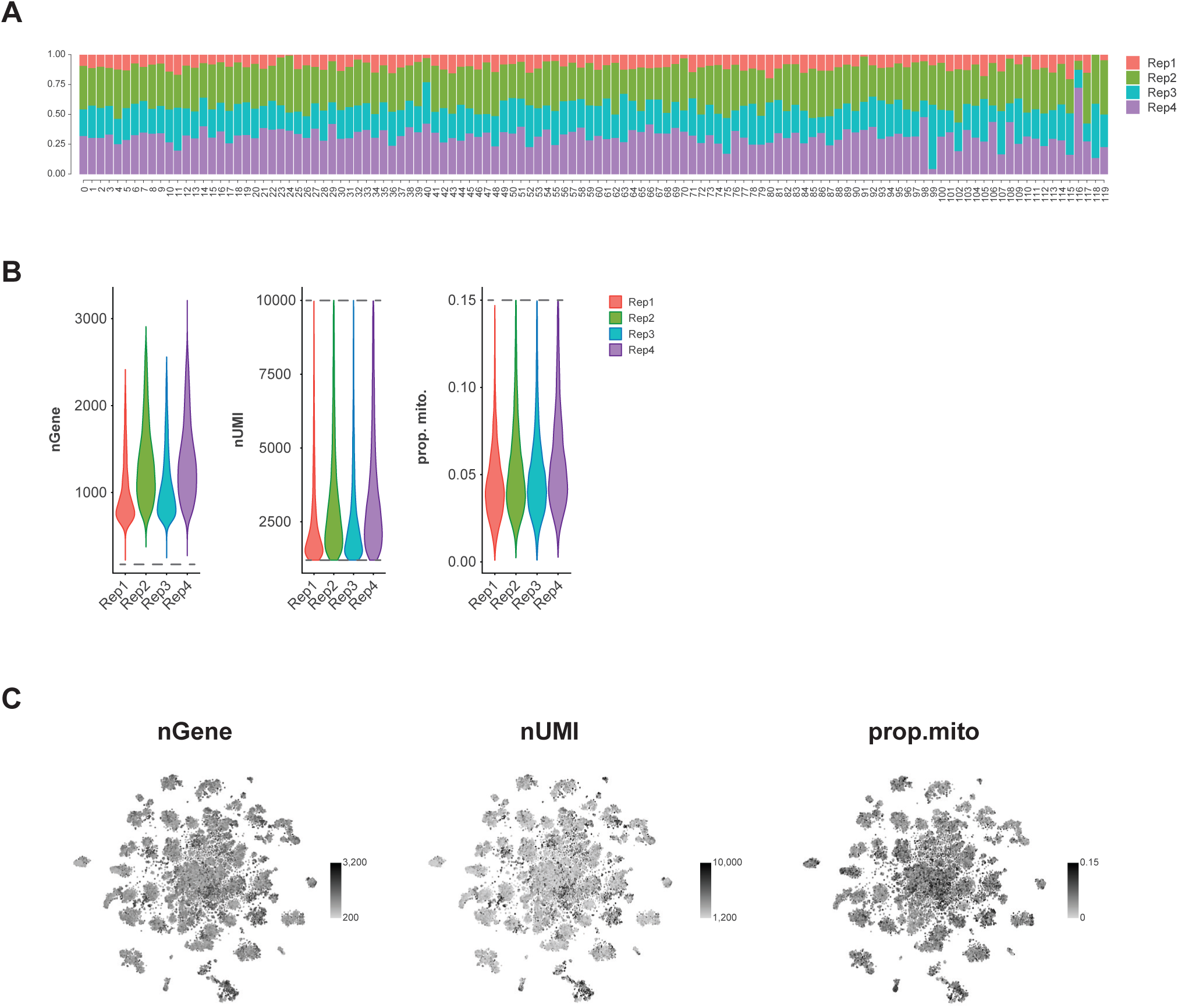
Replicate contributions and quality control. A. Proportional contribution of each replicate to each cluster in the t-SNE plot. B. Violin plots of the number of expressed genes (nGene), number of transcripts (nUMI) and proportion of transcripts that are mitochondrial (prop.mito) for each replicate. Imposed cut-offs are denoted with dashed lines. C. t-SNE plots showing the spatial distribution of the number of expressed genes (nGene), number of transcripts (nUMI) and proportion of transcripts that are mitochondrial (prop.mito).

**Figure 1-Figure supplement 4:**
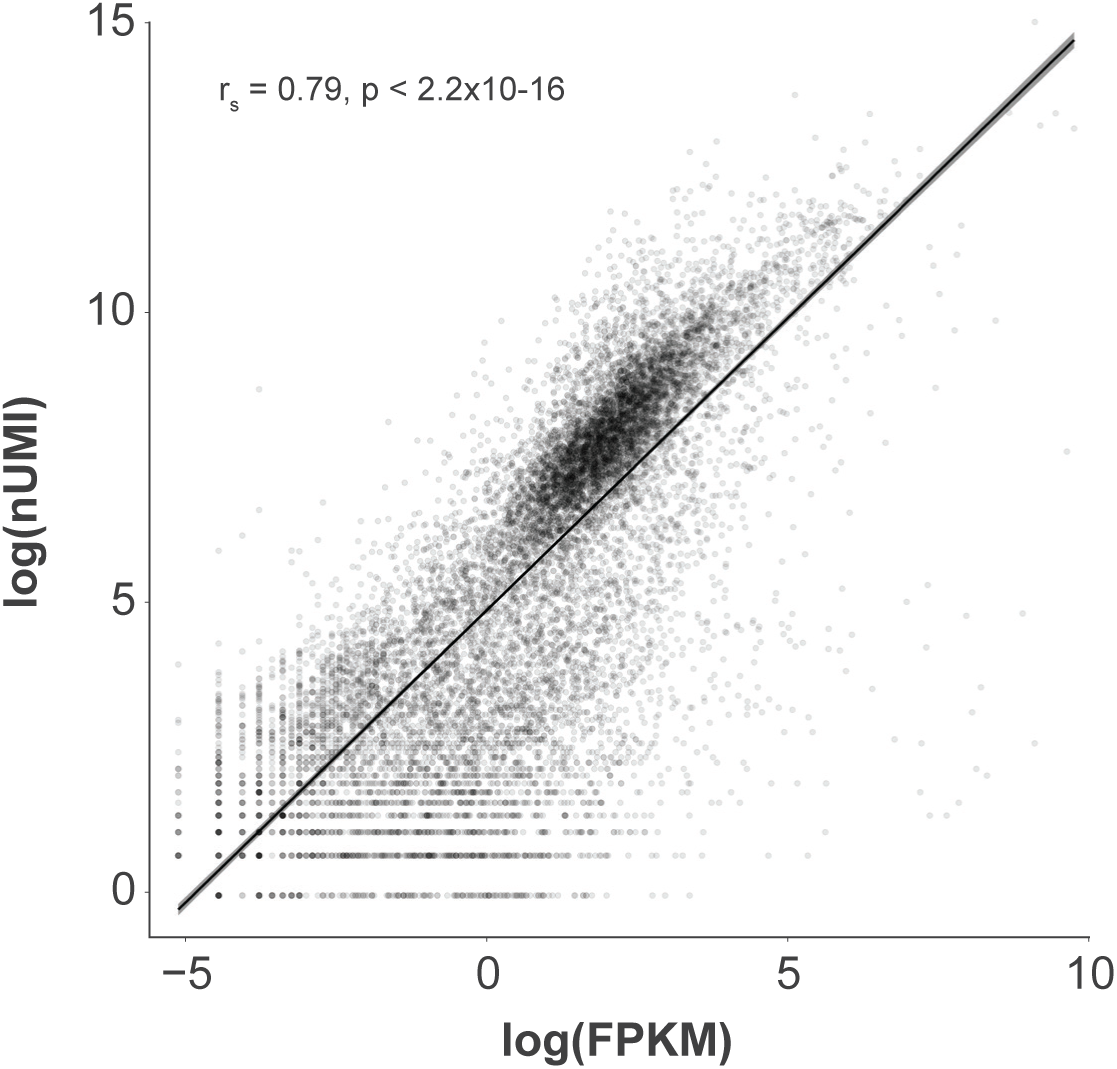
Single-cell vs. bulk RNA-seq expression in the VNC. Scatter plot showing the relationship between the expression level of each gene in this filtered single-cell data set (log(nUMI)) and the FlyAtlas 2 VNS data set (log(FPKM)).

**Figure 2-Figure supplement 1:**
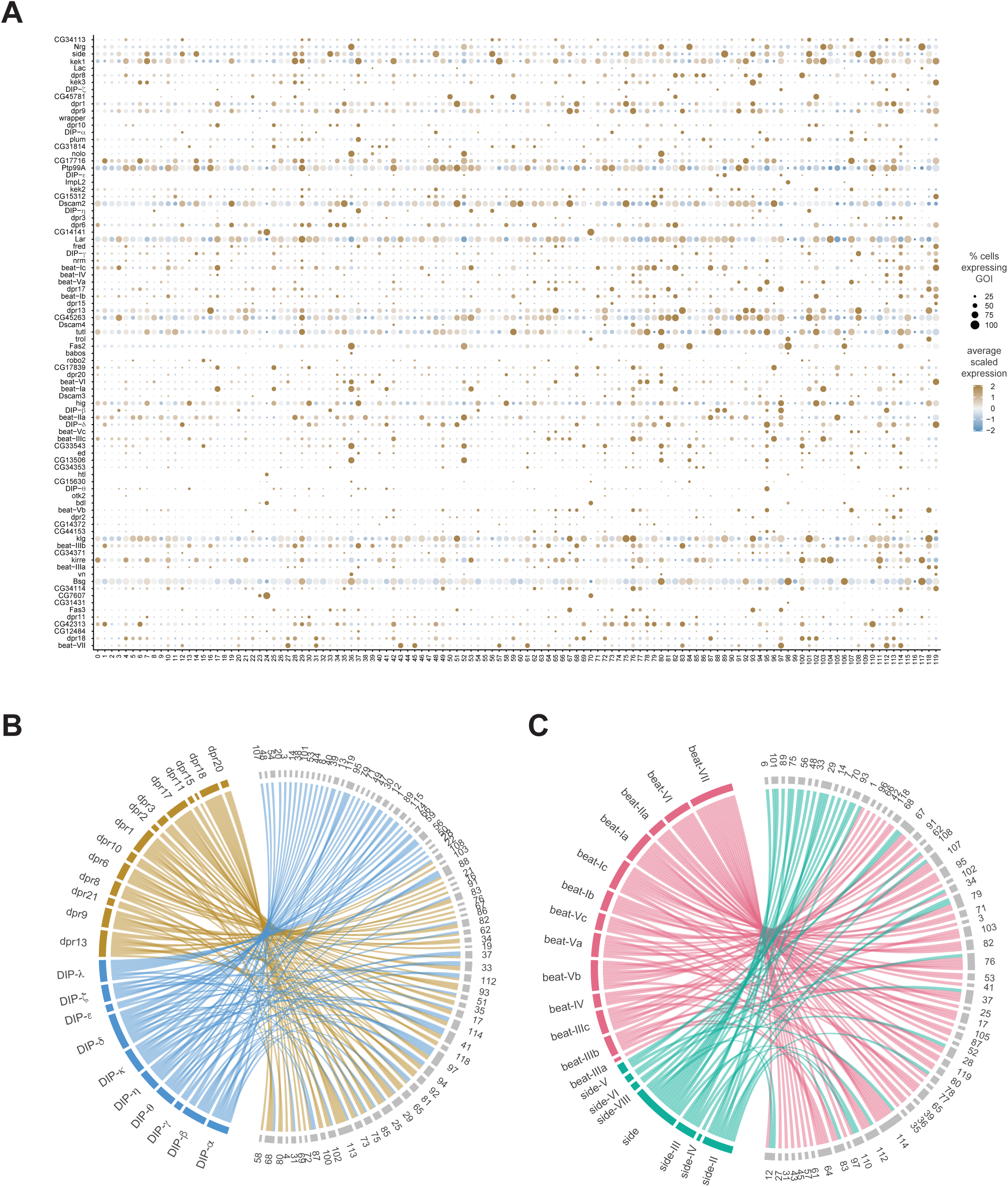
Relationships between IgSF subfamilies in defining cluster identity. A. Dot plot of the expression of IgSF cluster markers (left) across clusters (below). The gadient coloured fill represents the scaled, normalized expression level of each gene, and the radius of the dot represents the percent of cells expressing the gene of interest (GOI). B. Chord diagram showing dpr (left, brown) and DIP (left, blue) cluster markers with the clusters for which they are significantly enriched (right, in grey). C. Chord diagram showing Beat (left, green) and Side (left, pink) cluster markers with the clusters for which they are significantly enriched (right, in grey).

**Figure 2-Figure supplement 2:**
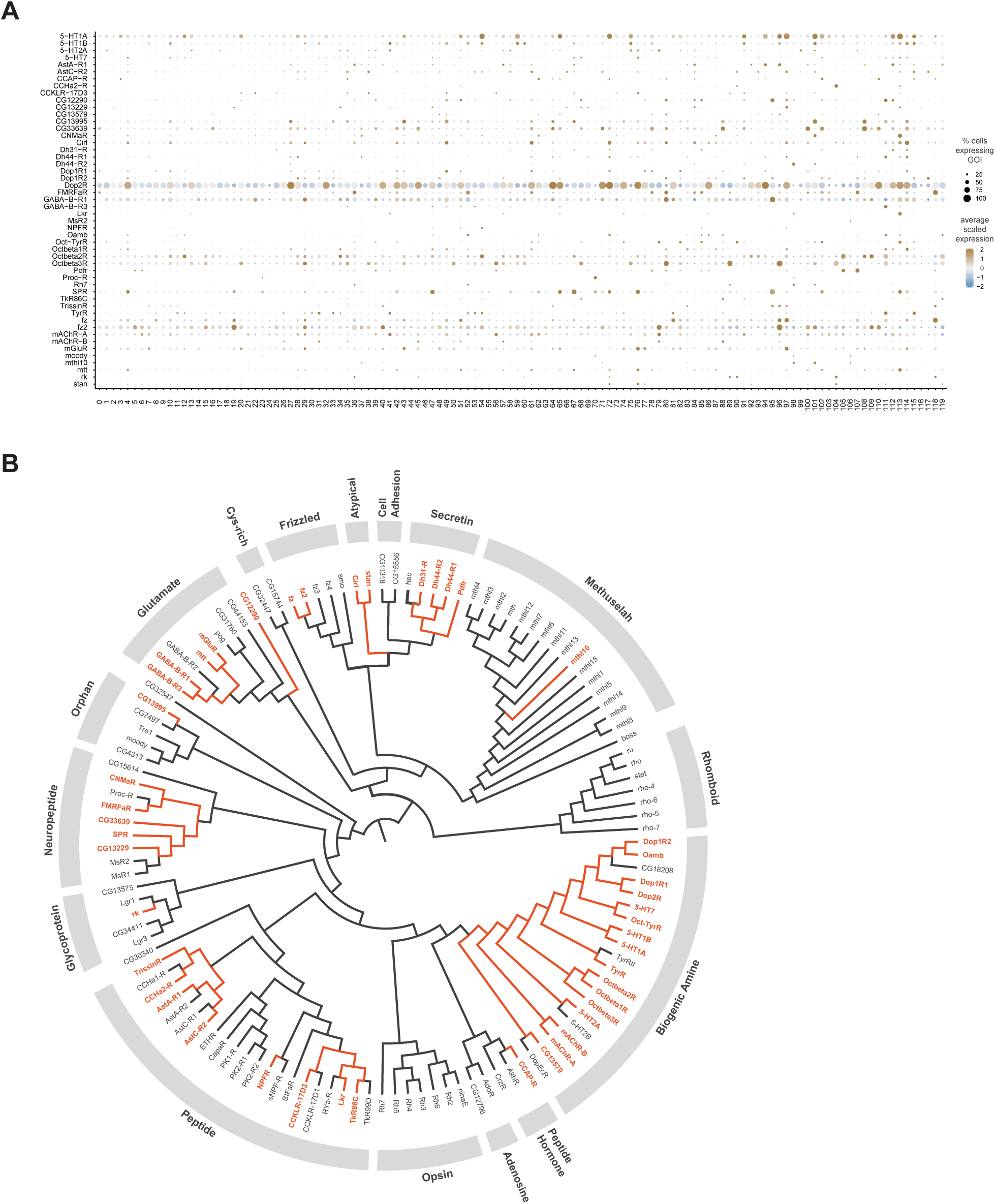
GPCR neuronal cluster markers. A. Dot plot of the expression of GPCR cluster markers(left) across clusters (below). The gadient coloured fill represents the scaled, normalized expression level of each gene, and the radius of the dot represents the percent of cells expressing the gene of interest (GOI). B. Phylogenetic tree of Drosophila GPCRs. GPCRs found as neuronal cluster markers are highlighted in orange. The multiple sequence alignment (not shown) and the phylogenetic tree were created with Clustal Omega and FigTreeV1.4.4, respectively (adapted from Hanlon and Andrew, 2015).

**Figure 2-Figure supplement 3:**
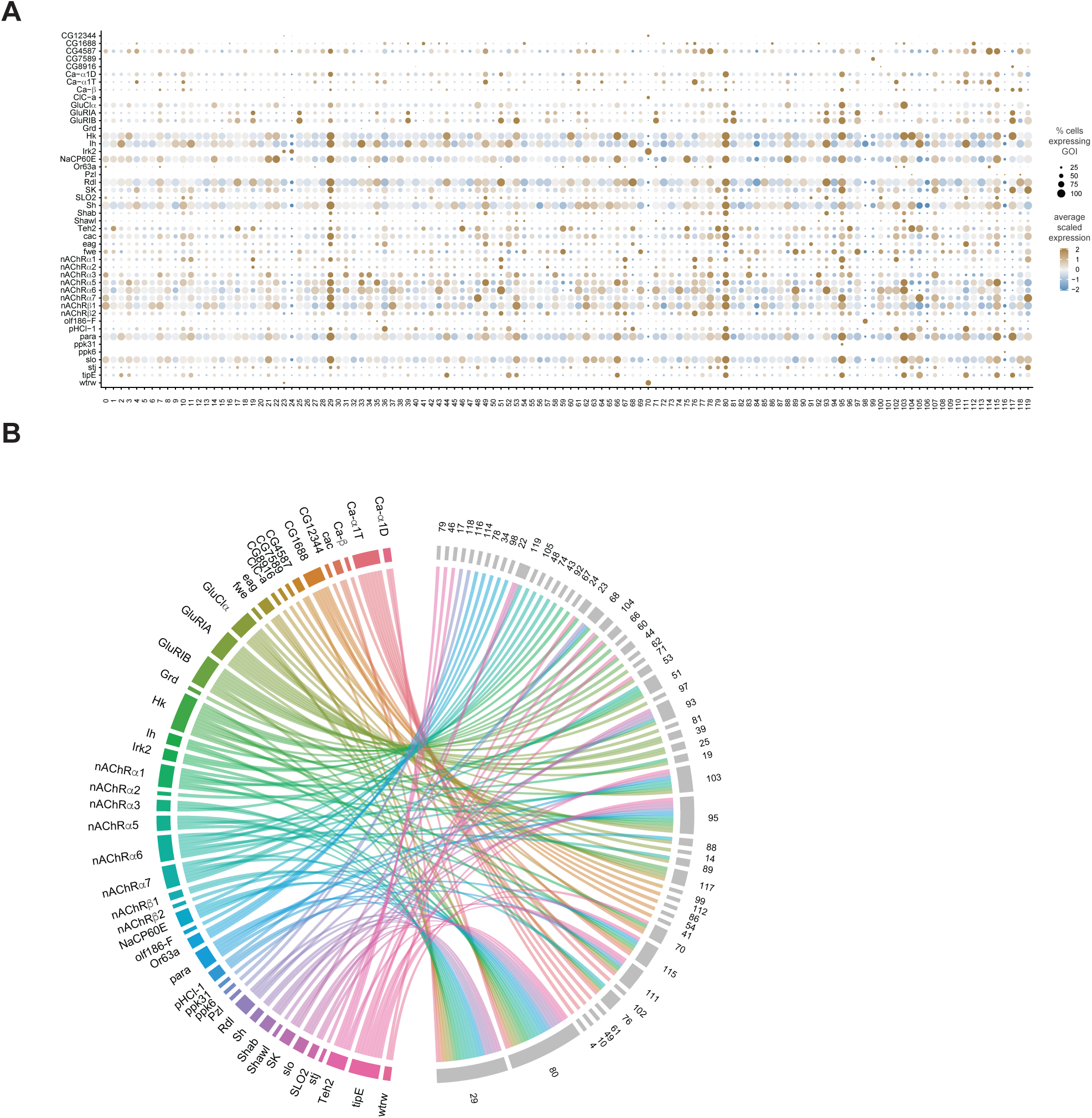
Ion channel neuronal cluster markers. A. Dot plot of the expression of ion channel cluster markers (left) across clusters (below). The gadient coloured fill represents the scaled, normalized expression level of each gene, and the radius of the dot represents the percent of cells expressing the gene of interest (GOI). B. Chord diagram illustrating the relationship between the ion channel cluster markers (left, colored) and the clusters (right, grey) for which they are significantly enriched.

**Figure 2-Figure supplement 4:**
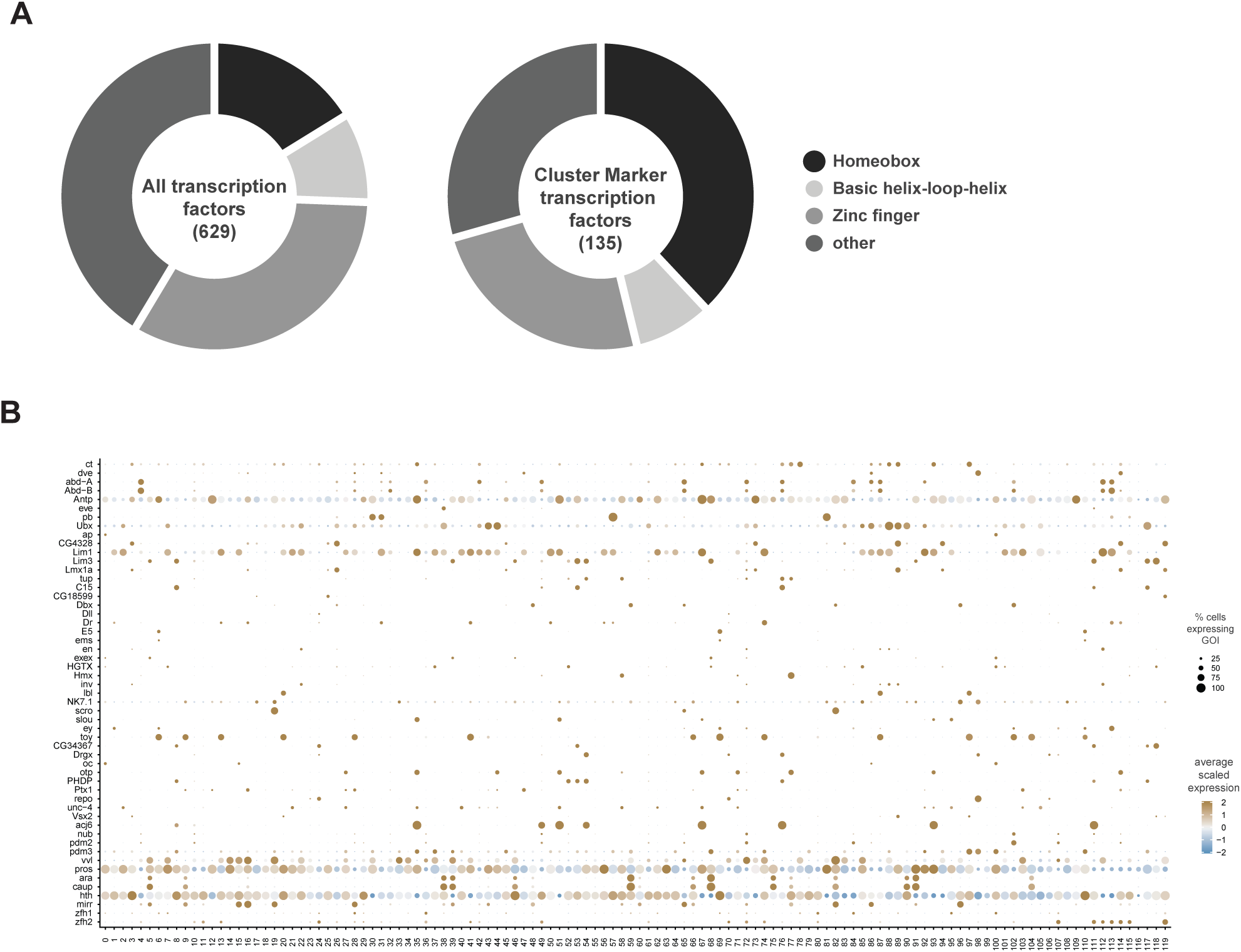
TF neuronal cluster markers. A. Doughnut graphs representing proportion of TF types found in the entire *Drosophila* genome (left) versus those found as neuronal cluster markers (right), based on Flybase Gene Group annotations. B. Dot plot of the expression of homeodomain TF cluster markers (left) across clusters (below). The gadient coloured fill represents the scaled, normalized expression level of each gene, and the radius of the dot represents the percent of cells expressing the gene of interest (GOI).

**Figure 4-Figure supplement 1:**
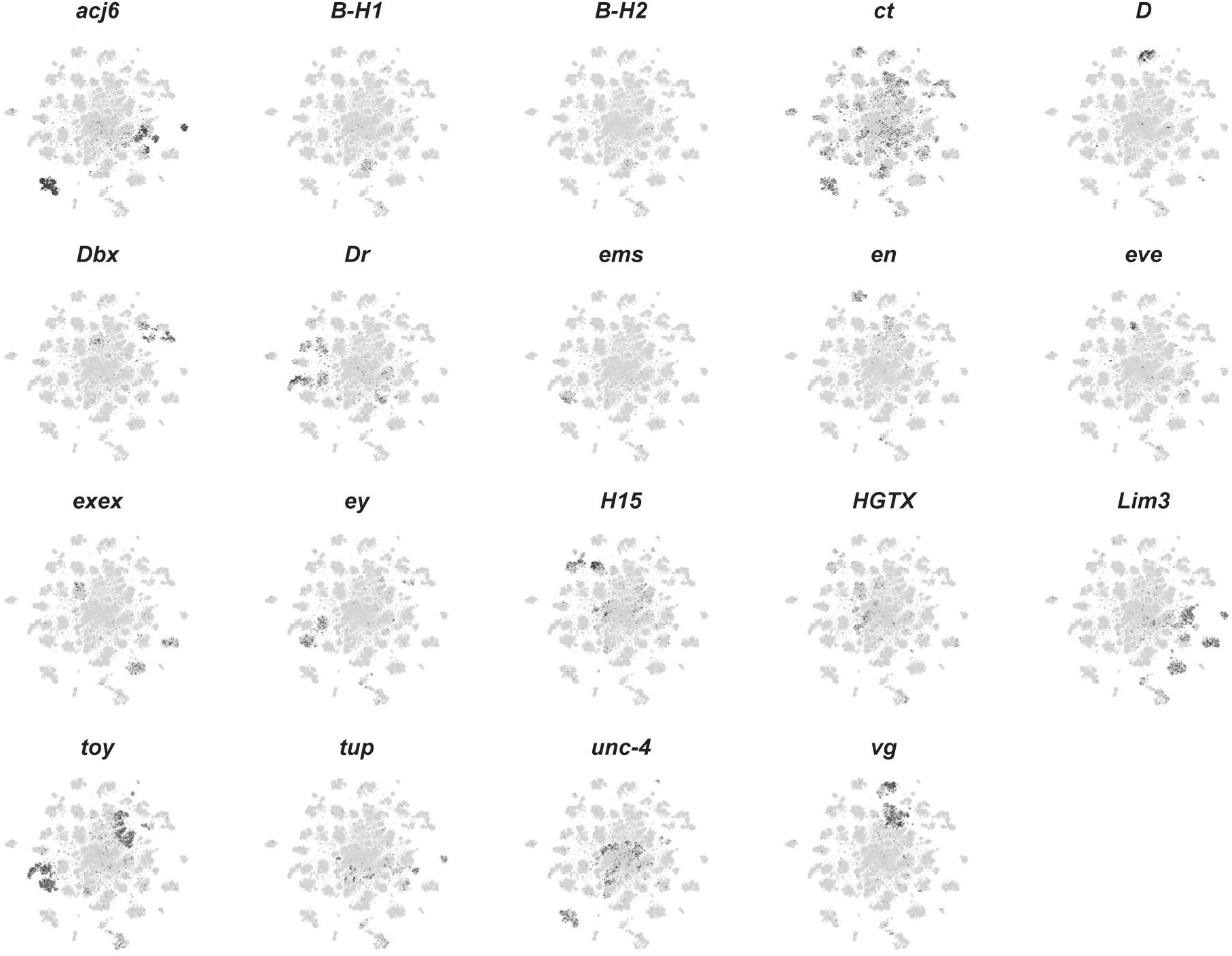
Neuroblast lineage marker gene expression in the VNC. A. t-SNE plots of the neuroblast lineage marker gene expression, shown in black, intensity is proportional to the log-normalized expression levels.

**Figure 4-Figure supplement 2:**
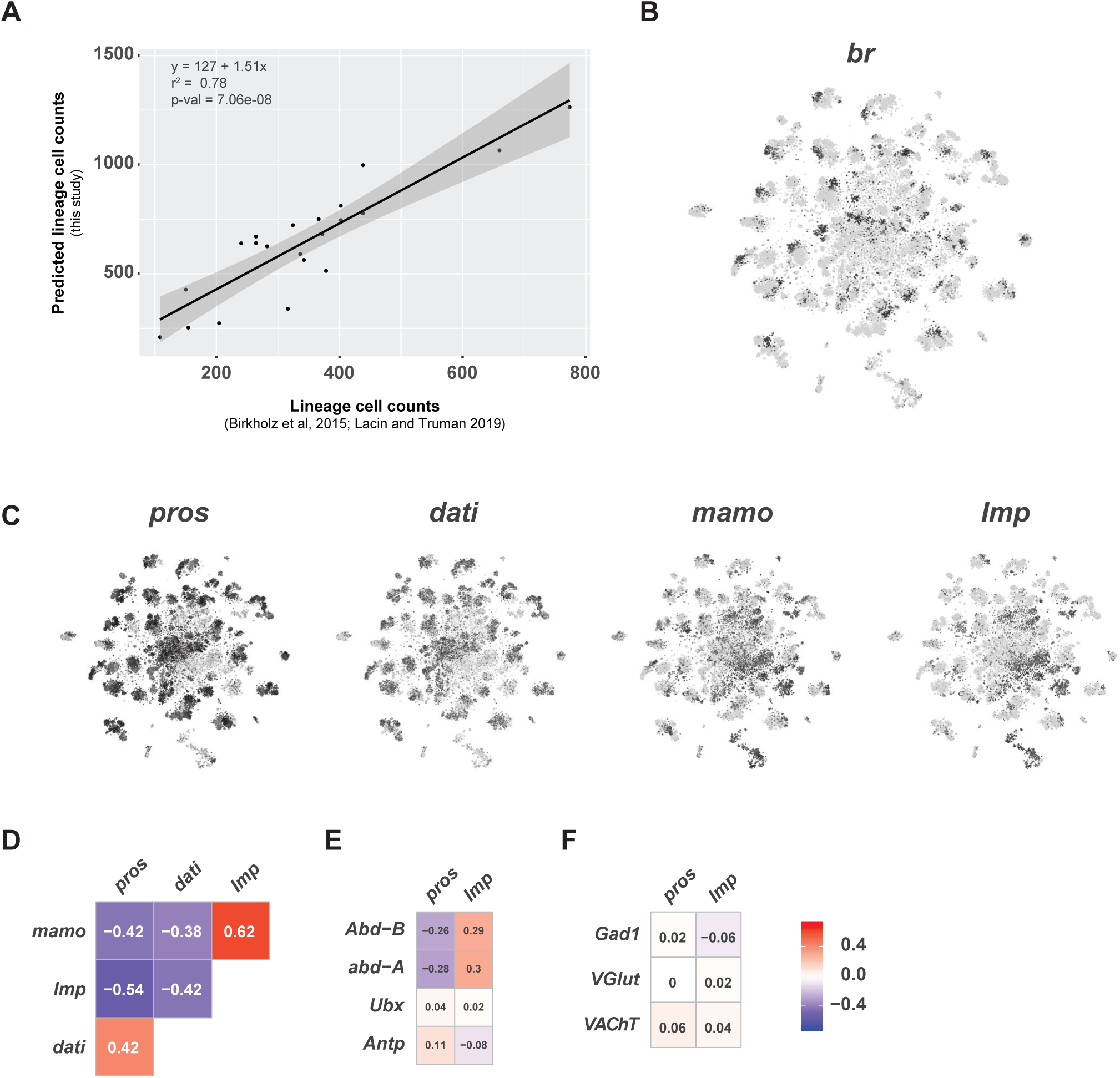
A. Scatter plot of hemilineage cell counts from thoracic neuromeres (calculated from Birkholz et al., 2015 and Lacin et al., 2019) compared to cell counts from predicted hemilineage identity in this single-cell data set (used pros+, abd-A-, Abd-B- cells to approximate postembryonic, thoracic hemilineages). B. t-SNE plot of br expression showing regional expression in clusters throughout the VNS. Expression shown in black, intensity is proportional to the log-normalized expression levels. C. t-SNE plots of pros, dati, mamo, and Imp in the VNC. Expression shown in black, intensity is proportional to the log-normalized expression levels. D. Heatmaps showing Pearson correlations coefficients (in text) of expression for *pros, dati, mamo, and Imp*. E. Heatmaps showing Pearson correlations coefficients (in text) of expression for *pros* and *Imp* with the Hox genes *Abd-B*, *abd-A*, *Ubx,* and *Antp*. F. Heatmaps showing Pearson correlations coefficients (in text) of expression for *pros* and *Imp* with fast-acting neurotransmitter markers *Gad1*, *VGlut,* and *VAChT*. Panels D-F share the same scale.

**Figure 5-Figure supplement 1:**
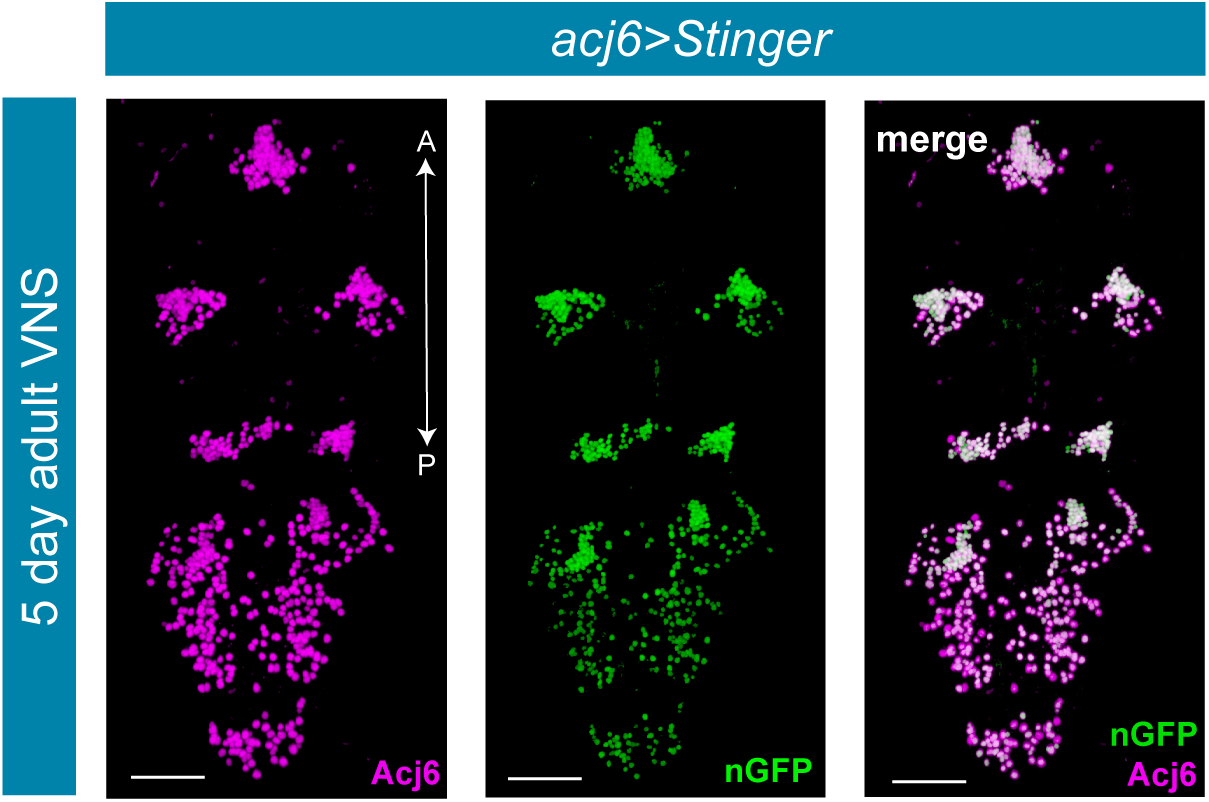
*acj6^GAL4^* recapitulates Acj6 protein expression in the adult VNC. Co-expression of *acj6^GAL4^* driven expression of *UAS-Stinger* (nGFP; green) and endogenous Acj6 protein expression (anti-Acj6, magenta) in the 5-day old adult VNC. Merged expression shown on right in white. A, Anterior, P, posterior; scale bars = 50 μm.

**Figure 5-Figure supplement 2:**
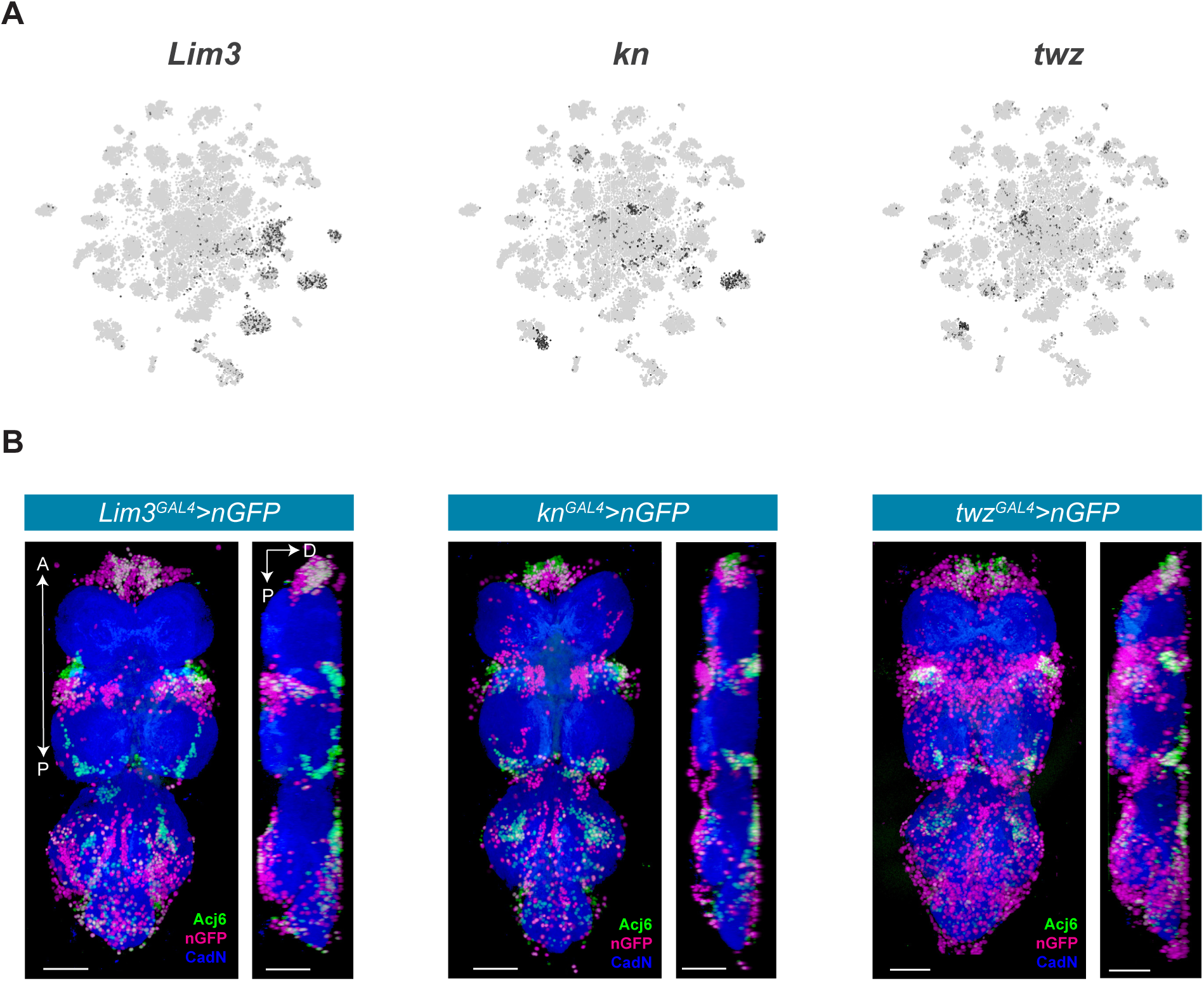
Novel lineage marker expression in the VNC. A. t-SNE plots of *Lim3* (A), *kn* (B) and *twz* (C) expression in the VNC. B. Co-expression of Acj6 (anti-Acj6; green) with *Lim3^GAL4^*, *kn^GAL4^,* and *twz^GAL4^* driven expression of *UAS-Stinger* (nGFP; magenta) in the 5-day adult VNC; coronal (left) and sagittal (right) views. A, Anterior, P, posterior, D, dorsal; Neuropil is counterstained (CadN; blue); scale bars = 50 μm.

**Figure 6-Figure supplement 1:**
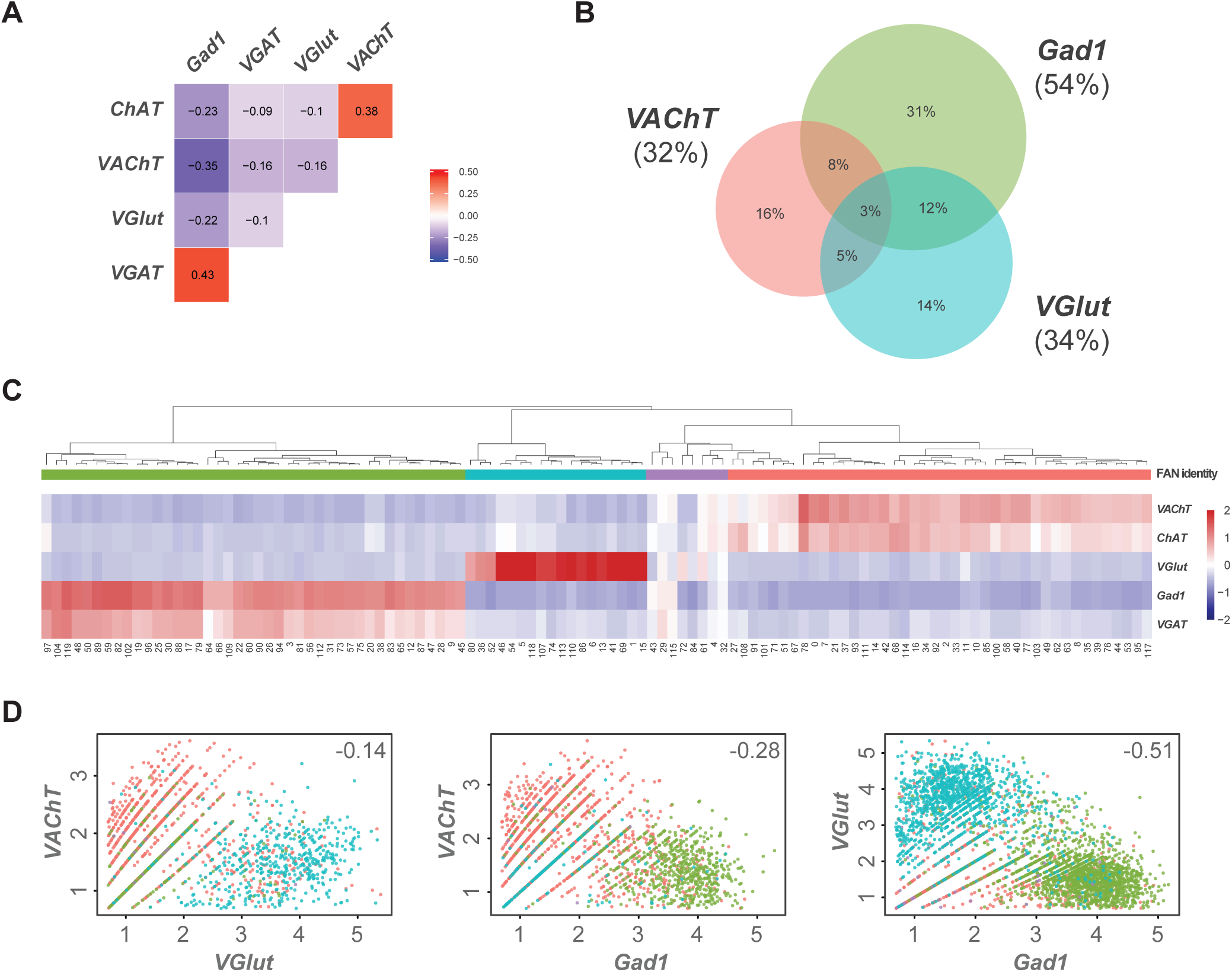
Expression of fast-acting neurotransmitter biomarkers in the VNC. A. Heatmap of Pearson correlation values for the expression fast-acting neurotransmitter biomarkers. B. Venn diagram showing the percent of cells with overlapping expression of *VAChT*, *VGlut* and *Gad1*. C. Heatmap of the mean scaled log-normalized expression, by cluster, of the fast-acting neurotransmitter biomarkers *VAChT/ChAT* (acetylcholine), *VGlut* (glutamate) and *VGAT/Gad1* (GABA). Cell clusters have been hierarchically grouped. Assigned FAN identity is color coded by bars below the dendrogram. D. Scatter plots showing the expression levels of fast-acting neurotransmitter biomarkers in pairwise co-expressing cells. Each dot represents a cell. Cells have been color coded by their assigned FAN identity. Pearson correlation coefficients are inset.

**Figure 7—source data 1: List of marker genes for the *Vmat*^+^ sub-clusters shown in** Figure 7C.

Table showing the average log-fold change values of monoaminergic cluster-discriminative marker genes, including adjusted p-values. pct.1 is the proportion of cells that express the gene in the cluster, pct.2 is the proportion of cells that express the gene in the other clusters.

**Figure 8-Figure supplement 1:**
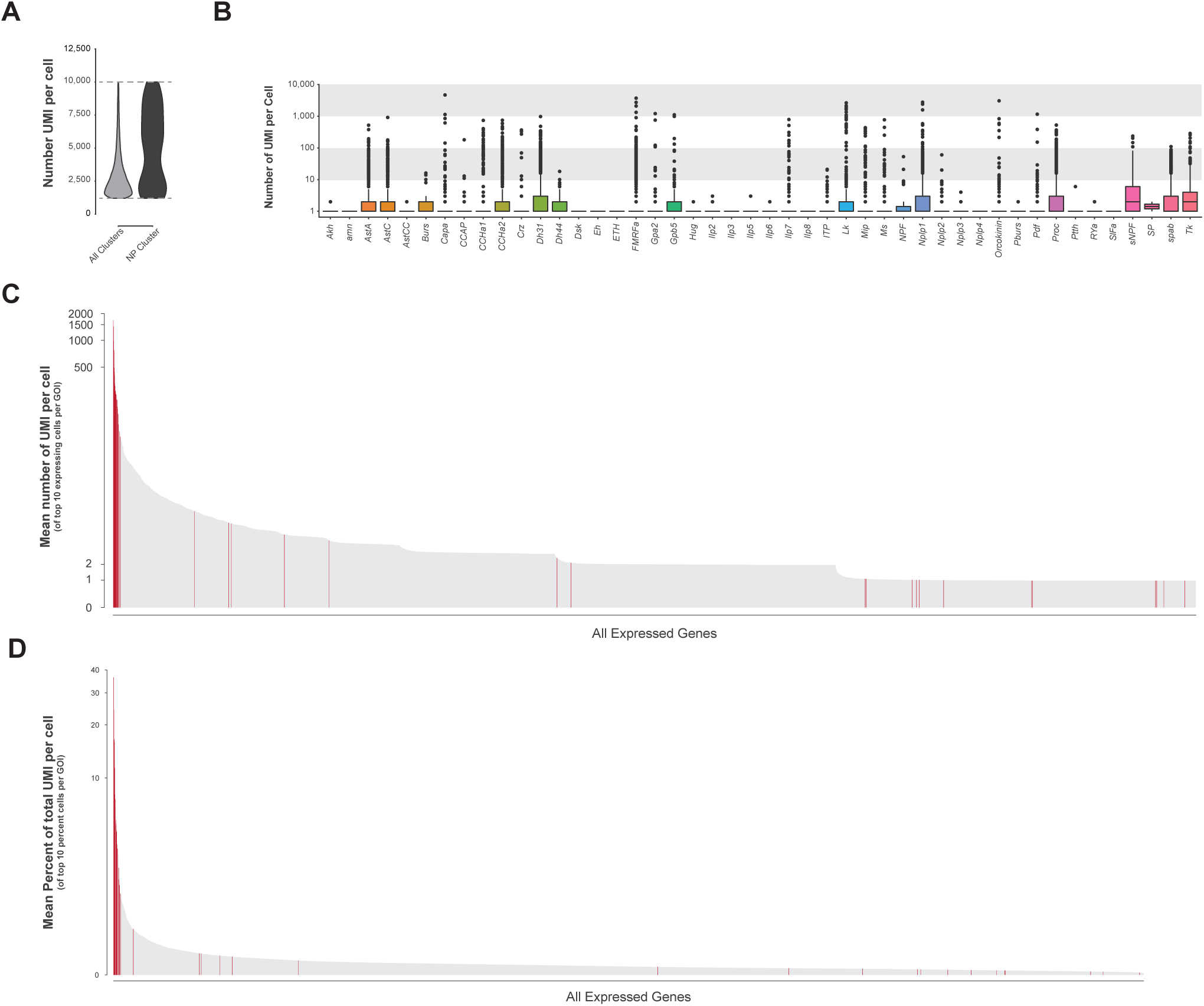
Neuropeptide gene expression levels. A. Violin plot showing the number of transcripts (UMI) per cell in the whole VNC data set (all clusters) compared to neuropeptide-expressing cluster 84. Data represents expression between 1,200-10,000 UMI (dashed lines). B. Box plots representing number of UMI per cell of all expressed neuropeptide genes across the whole VNC data set. C. Ranked mean number of UMI per cell of the top 10 expressing cells for each GOI. Neuropeptide genes are highlighted in red, many of which have the highest number of UMI/cell in the genome. D. Ranked mean percent of UMI per cell (GOI UMI/total UMI) of the top 10 proportion of UMI cells for each GOI. Neuropeptide genes are highlighted in red, many of which have the highest proportion of transcripts per cell in the genome.

**Figure 8-Figure supplement 2:**
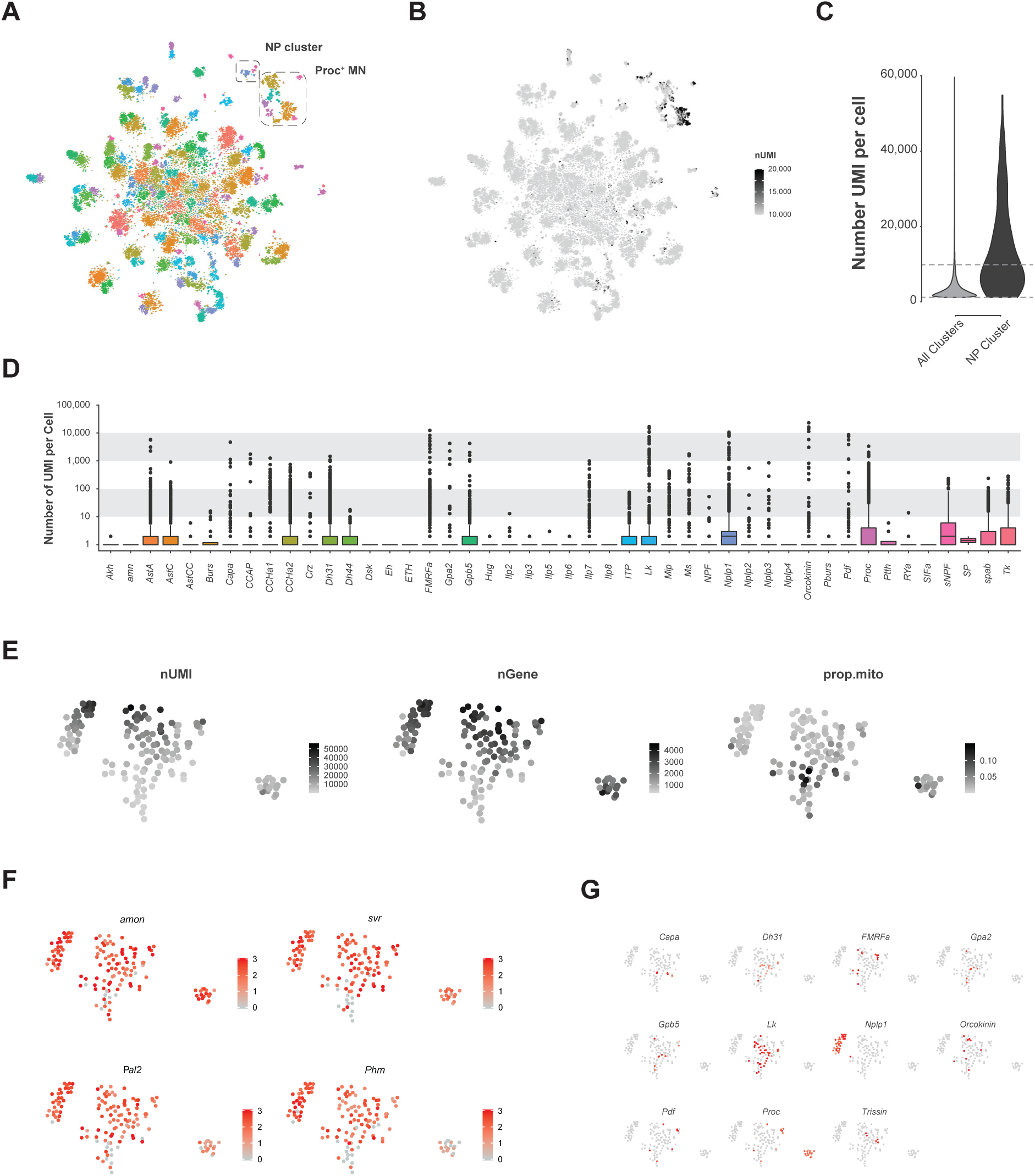
Analysis of VNC data removing upper limit on transcript number (UMI) per cell. **A.** Re-analyzed t-SNE plot of VNC data without the 10,000 UMI high cut-off. Neuropeptide (NP*) enriched cluster and Proctolin expressing motor neuron (Proc+ MN) cluster are highlighted. **B.** t-SNE of the number of UMI. Color is scaled from 10,000 to 20,000 UMI. Most cells with greater than 10,000 UMI are in the NP cluster or the *Proc^+^* MN cluster. **C.** Violin plot showing the number of UMI per cell without 10,000 UMI filtering cut-off in the whole VNS data set (all clusters) compared to the NP cluster. Dashed lines represent the low cut-off of 1,200 UMI and the now removed cut-off of 10,000 UMI. **D.** Box plots representing number of UMI per cell of all expressed neuropeptide genes across the whole VNS data set without the 10,000 UMI filtering cut-off. Multiple NP genes exceed the expression level of 10,000 UMI. **E.** t-SNE plots of the NP cluster showing the spatial distribution of the number of expressed genes (nGene), number of transcripts (nUMI) and proportion of transcripts that are mitochondrial (prop.mito). F, G Expression of neuropeptide processing enzymes (F) and multiple neuropeptide genes (G) in t-SNE of NP cluster. Intensity of red is proportional to the log-normalized expression level

**Figure 8-Figure supplement 3:**
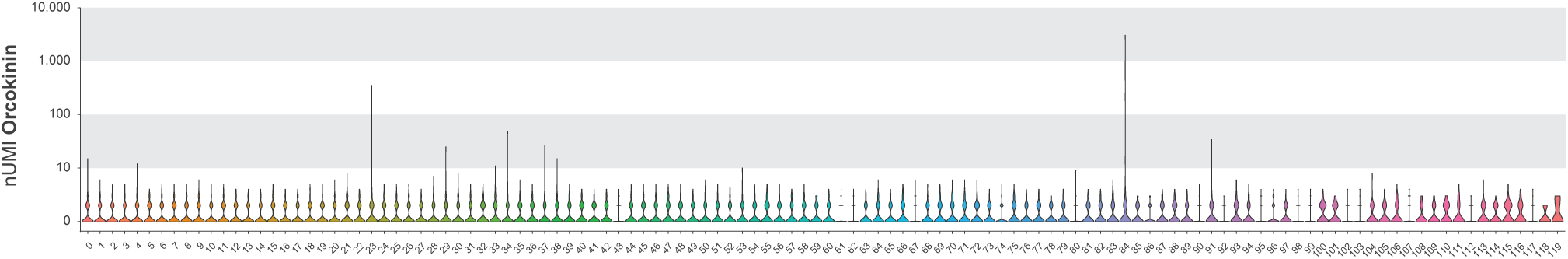
*Orcokinin* expression analysis. Violin plot of raw UMI *Orcrokinin* expression by cluster.

**Figure 9—source data 1: List of marker genes for the glial sub-clusters shown in** Figure 9B.

Table showing the average log-fold change values of glial cluster-discriminative marker genes, including adjusted p-values. pct.1 is the proportion of cells that express the gene in the cluster, pct.2 is the proportion of cells that express the gene in the other clusters.

**Figure 9 supplement 1:**
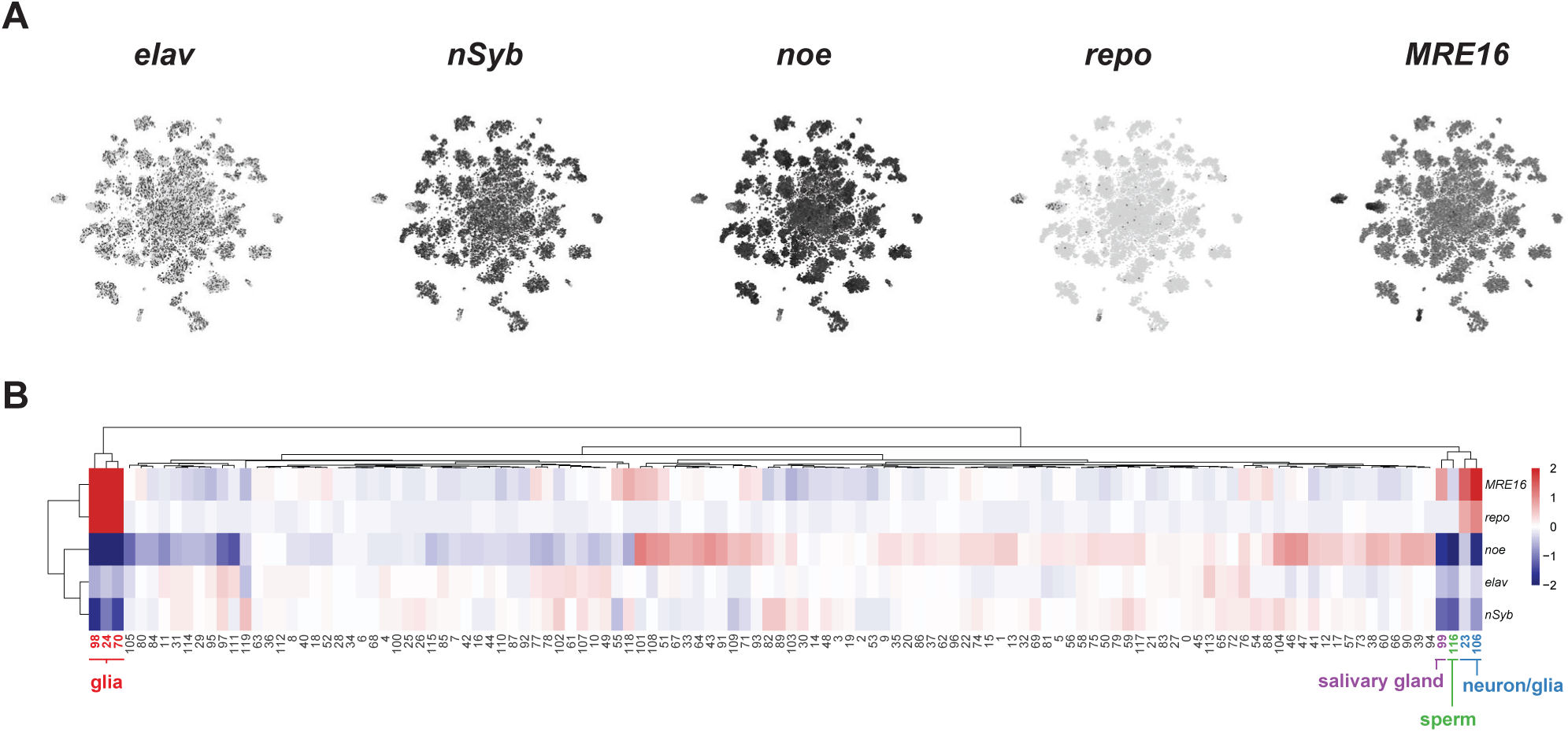
Identification of neuronal and glia clusters. A. t-SNE plots showing distribution of cells expressing known neuronal (*elav, nSyb, noe*) and glial (*repo, MRE16*) marker genes. B. Heatmap of the mean scaled log-normalized expression, by cluster, of glial and neuronal marker genes. Glial cluster numbers are shown in red, salivary gland in purple, sperm in green and neuron/glia mixed cluster numbers in blue.

## References

Agrawal, T., Sadaf, S., and Hasan, G. (2013). A genetic RNAi screen for IP(3)/Ca(2)(+) coupled GPCRs in *Drosophila* identifies the PdfR as a regulator of insect flight. PLoS Genet 9, e1003849.

Anderson, M.G., Perkins, G.L., Chittick, P., Shrigley, R.J., and Johnson, W.A. (1995). *drifter*, a *Drosophila* POU-domain transcription factor, is required for correct differentiation and migration of tracheal cells and midline glia. Genes Dev 9, 123–137.

Angelini, D.R., and Kaufman, T.C. (2005). Insect appendages and comparative ontogenetics. Dev Biol 286, 57–77.

Baek, M., Enriquez, J., and Mann, R.S. (2013). Dual role for Hox genes and Hox co-factors in conferring leg motoneuron survival and identity in *Drosophila*. Development 140, 2027–2038.

Baek, M., and Mann, R.S. (2009). Lineage and birth date specify motor neuron targeting and dendritic architecture in adult *Drosophila*. J Neurosci 29, 6904–6916.

Barolo, S., Carver, L.A., and Posakony, J.W. (2000). GFP and beta-galactosidase transformation vectors for promoter/enhancer analysis in *Drosophila*. Biotechniques 29, 726, 728, 730, 732.

Bates, A.S., Janssens, J., Jefferis, G.S., and Aerts, S. (2019). Neuronal cell types in the fly: single-cell anatomy meets single-cell genomics. Curr Opin Neurobiol 56, 125–134.

Bidaye, S.S., Machacek, C., Wu, Y., and Dickson, B.J. (2014). Neuronal control of *Drosophila* walking direction. Science 344, 97–101.

Billeter, J.C., Villella, A., Allendorfer, J.B., Dornan, A.J., Richardson, M., Gailey, D.A., and Goodwin, S.F. (2006). Isoform-specific control of male neuronal differentiation and behavior in *Drosophila* by the *fruitless* gene. Curr Biol 16, 1063–1076.

Birkholz, O., Rickert, C., Nowak, J., Coban, I.C., and Technau, G.M. (2015). Bridging the gap between postembryonic cell lineages and identified embryonic neuroblasts in the ventral nerve cord of *Drosophila melanogaster*. Biol Open 4, 420–434.

Bossing, T., Udolph, G., Doe, C.Q., and Technau, G.M. (1996). The embryonic central nervous system lineages of *Drosophila melanogaster*. I. Neuroblast lineages derived from the ventral half of the neuroectoderm. Dev Biol 179, 41–64.

Buchner, E., Buchner, S., Burg, M.G., Hofbauer, A., Pak, W.L., and Pollack, I. (1993). Histamine is a major mechanosensory neurotransmitter candidate in *Drosophila melanogaster*. Cell Tissue Res 273, 119–125.

Bürglin, T.R., and Affolter, M. (2016). Homeodomain proteins: an update. Chromosoma 125, 497–521.

Burrows, M. (1992). Local circuits for the control of leg movements in an insect. Trends Neurosci 15, 226–232.

Card, G., and Dickinson, M.H. (2008). Visually mediated motor planning in the escape response of *Drosophila*. Curr Biol 18, 1300–1307.

Carrillo, R.A., Ozkan, E., Menon, K.P., Nagarkar-Jaiswal, S., Lee, P.T., Jeon, M., Birnbaum, M.E., Bellen, H.J., Garcia, K.C., and Zinn, K. (2015). Control of Synaptic Connectivity by a Network of *Drosophila* IgSF Cell Surface Proteins. Cell 163, 1770–1782.

Cavodeassi, F., Modolell, J., and Gomez-Skarmeta, J.L. (2001). The Iroquois family of genes: from body building to neural patterning. Development 128, 2847–2855.

Chen, C.L., Hermans, L., Viswanathan, M.C., Fortun, D., Aymanns, F., Unser, M., Cammarato, A., Dickinson, M.H., and Ramdya, P. (2018). Imaging neural activity in the ventral nerve cord of behaving adult *Drosophila*. Nat Commun 9, 4390.

Chen, J., Choi, M.S., Mizoguchi, A., Veenstra, J.A., Kang, K., Kim, Y.J., and Kwon, J.Y. (2015). Isoform-specific expression of the neuropeptide orcokinin in Drosophila melanogaster. Peptides 68, 50–57.

Chintapalli, V.R., Wang, J., Herzyk, P., Davies, S.A., and Dow, J.A. (2013). Data-mining the FlyAtlas online resource to identify core functional motifs across transporting epithelia. BMC Genomics 14, 518.

Clyne, J.D., and Miesenbock, G. (2008). Sex-specific control and tuning of the pattern generator for courtship song in *Drosophila*. Cell 133, 354–363.

Cognigni, P., Felsenberg, J., and Waddell, S. (2018). Do the right thing: neural network mechanisms of memory formation, expression and update in *Drosophila*. Curr Opin Neurobiol 49, 51–58.

Cole, S.H., Carney, G.E., McClung, C.A., Willard, S.S., Taylor, B.J., and Hirsh, J. (2005). Two functional but noncomplementing *Drosophila* tyrosine decarboxylase genes: distinct roles for neural tyramine and octopamine in female fertility. J Biol Chem 280, 14948–14955.

Court, R.C., Armstrong, J., Börner, J., Card, G., Costa, M., Dickinson, M., Duch, C., Korff, W., Mann, R., Merritt, D., et al. (2017). A Systematic Nomenclature for the Drosophila Ventral Nervous System. bioRxiv, 122952.

Crickmore, M.A., and Vosshall, L.B. (2013). Opposing dopaminergic and GABAergic neurons control the duration and persistence of copulation in Drosophila. Cell 155, 881–893.

Croset, V., Treiber, C.D., and Waddell, S. (2018). Cellular diversity in the *Drosophila* midbrain revealed by single-cell transcriptomics. Elife 7.

Crozatier, M., Valle, D., Dubois, L., Ibnsouda, S., and Vincent, A. (1996). Collier, a novel regulator of *Drosophila* head development, is expressed in a single mitotic domain. Curr Biol 6, 707–718.

Cully, D.F., Paress, P.S., Liu, K.K., Schaeffer, J.M., and Arena, J.P. (1996). Identification of a *Drosophila melanogaster* glutamate-gated chloride channel sensitive to the antiparasitic agent avermectin. J Biol Chem 271, 20187–20191.

Dason, J.S., Romero-Pozuelo, J., Marin, L., Iyengar, B.G., Klose, M.K., Ferrus, A., and Atwood, H.L. (2009). Frequenin/NCS-1 and the Ca2+-channel alpha1-subunit co-regulate synaptic transmission and nerve-terminal growth. J Cell Sci 122, 4109–4121.

Davie, K., Janssens, J., Koldere, D., De Waegeneer, M., Pech, U., Kreft, L., Aibar, S., Makhzami, S., Christiaens, V., Bravo Gonzalez-Blas, C., et al. (2018). A Single-Cell Transcriptome Atlas of the Aging *Drosophila* Brain. Cell 174, 982–998 e920.

Deneris, E.S., and Hobert, O. (2014). Maintenance of postmitotic neuronal cell identity. Nat Neurosci 17, 899–907.

DeSalvo, M.K., Hindle, S.J., Rusan, Z.M., Orng, S., Eddison, M., Halliwill, K., and Bainton, R.J. (2014). The *Drosophila* surface glia transcriptome: evolutionary conserved blood-brain barrier processes. Front Neurosci 8, 346.

DiAntonio, A., Burgess, R.W., Chin, A.C., Deitcher, D.L., Scheller, R.H., and Schwarz, T.L. (1993). Identification and characterization of *Drosophila* genes for synaptic vesicle proteins. J Neurosci 13, 4924–4935.

Doherty, J., Logan, M.A., Tasdemir, O.E., and Freeman, M.R. (2009). Ensheathing glia function as phagocytes in the adult *Drosophila* brain. J Neurosci 29, 4768–4781.

Enriquez, J., Rio, L.Q., Blazeski, R., Bellemin, S., Godement, P., Mason, C., and Mann, R.S. (2018). Differing Strategies Despite Shared Lineages of Motor Neurons and Glia to Achieve Robust Development of an Adult Neuropil in *Drosophila*. Neuron 97, 538–554 e535.

Estacio-Gómez, A., Hassan, A., Walmsley, E., Lee, L., and Southall, T.D. (2019). Dynamic neurotransmitter specific transcription factor expression profiles during Drosophila development. bioRxiv, 830315.

Etheredge, J. (2017). Transcriptional profiling of *Drosophila* larval ventral nervous system hemilineages. In Physiology, Development, and Neuroscience (University of Cambridge).

Gowda, S.B.M., Paranjpe, P.D., Reddy, O.V., Thiagarajan, D., Palliyil, S., Reichert, H., and VijayRaghavan, K. (2018). GABAergic inhibition of leg motoneurons is required for normal walking behavior in freely moving *Drosophila*. Proc Natl Acad Sci U S A 115, E2115–E2124.

Greer, C.L., Grygoruk, A., Patton, D.E., Ley, B., Romero-Calderon, R., Chang, H.Y., Houshyar, R., Bainton, R.J., Diantonio, A., and Krantz, D.E. (2005). A splice variant of the *Drosophila* vesicular monoamine transporter contains a conserved trafficking domain and functions in the storage of dopamine, serotonin, and octopamine. J Neurobiol 64, 239–258.

Grillner, S. (2006). Biological pattern generation: the cellular and computational logic of networks in motion. Neuron 52, 751–766.

Gu, Z., Gu, L., Eils, R., Schlesner, M., and Brors, B. (2014). circlize Implements and enhances circular visualization in R. Bioinformatics 30, 2811–2812.

Hamanaka, Y., Park, D., Yin, P., Annangudi, S.P., Edwards, T.N., Sweedler, J., Meinertzhagen, I.A., and Taghert, P.H. (2010). Transcriptional orchestration of the regulated secretory pathway in neurons by the bHLH protein DIMM. Curr Biol 20, 9–18.

Hanlon, C.D., and Andrew, D.J. (2015). Outside-in signaling--a brief review of GPCR signaling with a focus on the *Drosophila* GPCR family. J Cell Sci 128, 3533–3542.

Hardie, R.C. (1987). Is histamine a neurotransmitter in insect photoreceptors? J Comp Physiol A 161, 201–213.

Harris, R.M., Pfeiffer, B.D., Rubin, G.M., and Truman, J.W. (2015). Neuron hemilineages provide the functional ground plan for the *Drosophila* ventral nervous system. Elife 4.

Hartenstein, V. (2018). Development of the Nervous System of Invertebrates. In The Oxford Handbook of Invertebrate Neurobiology, J.H. Byrne, ed. (Oxford University Press).

Hartenstein, V., Cruz, L., Lovick, J.K., and Guo, M. (2017). Developmental analysis of the dopamine-containing neurons of the *Drosophila* brain. J Comp Neurol 525, 363–379.

Hewes, R.S., Park, D., Gauthier, S.A., Schaefer, A.M., and Taghert, P.H. (2003). The bHLH protein Dimmed controls neuroendocrine cell differentiation in *Drosophila*. Development 130, 1771–1781.

Hobert, O., and Kratsios, P. (2019). Neuronal identity control by terminal selectors in worms, flies, and chordates. Curr Opin Neurobiol 56, 97–105.

Honegger, B., Galic, M., Kohler, K., Wittwer, F., Brogiolo, W., Hafen, E., and Stocker, H. (2008). Imp-L2, a putative homolog of vertebrate IGF-binding protein 7, counteracts insulin signaling in *Drosophila* and is essential for starvation resistance. J Biol 7, 10.

Howard, C.E., Chen, C.-L., Tabachnik, T., Hormigo, R., Ramdya, P., and Mann, R.S. (2019). Serotonergic modulation of walking in Drosophila. bioRxiv, 753624.

Huang, D.W., Sherman, B.T., and Lempicki, R.A. (2009). Systematic and integrative analysis of large gene lists using DAVID bioinformatics resources. Nat Protoc 4, 44–57.

Huang, Y., Ng, F.S., and Jackson, F.R. (2015). Comparison of larval and adult *Drosophila* astrocytes reveals stage-specific gene expression profiles. G3 (Bethesda) 5, 551–558.

Ito, K., Shinomiya, K., Ito, M., Armstrong, J.D., Boyan, G., Hartenstein, V., Harzsch, S., Heisenberg, M., Homberg, U., Jenett, A., et al. (2014). A systematic nomenclature for the insect brain. Neuron 81, 755–765.

Jois, S., Chan, Y.B., Fernandez, M.P., and Leung, A.K. (2018). Characterization of the Sexually Dimorphic *fruitless* Neurons That Regulate Copulation Duration. Front Physiol 9, 780.

Jonsson, T., Kravitz, E.A., and Heinrich, R. (2011). Sound production during agonistic behavior of male *Drosophila melanogaster*. Fly (Austin) 5, 29–38.

Joseph, R.M., and Carlson, J.R. (2015). *Drosophila* Chemoreceptors: A Molecular Interface Between the Chemical World and the Brain. Trends Genet 31, 683–695.

Kerner, P., Ikmi, A., Coen, D., and Vervoort, M. (2009). Evolutionary history of the *iroquois/Irx* genes in metazoans. BMC Evol Biol 9, 74.

Kikuta, S., Hagiwara-Komoda, Y., Noda, H., and Kikawada, T. (2012). A novel member of the trehalose transporter family functions as an H(+)-dependent trehalose transporter in the reabsorption of trehalose in malpighian tubules. Front Physiol 3, 290.

Kim, B., Shortridge, R.D., Seong, C., Oh, Y., Baek, K., and Yoon, J. (1998). Molecular characterization of a novel *Drosophila* gene which is expressed in the central nervous system. Mol Cells 8, 750–757.

Kim, H., Kirkhart, C., and Scott, K. (2017). Long-range projection neurons in the taste circuit of *Drosophila*. Elife 6.

Kimura, K., Sato, C., Koganezawa, M., and Yamamoto, D. (2015). *Drosophila* ovipositor extension in mating behavior and egg deposition involves distinct sets of brain interneurons. PLoS One 10, e0126445.

Knauf, F., Rogina, B., Jiang, Z., Aronson, P.S., and Helfand, S.L. (2002). Functional characterization and immunolocalization of the transporter encoded by the life-extending gene Indy. Proc Natl Acad Sci U S A 99, 14315–14319.

Konstantinides, N., Kapuralin, K., Fadil, C., Barboza, L., Satija, R., and Desplan, C. (2018). Phenotypic Convergence: Distinct Transcription Factors Regulate Common Terminal Features. Cell 174, 622–635 e613.

Lacin, H., Chen, H.M., Long, X., Singer, R.H., Lee, T., and Truman, J.W. (2019). Neurotransmitter identity is acquired in a lineage-restricted manner in the *Drosophila* CNS. Elife 8.

Lacin, H., and Truman, J.W. (2016). Lineage mapping identifies molecular and architectural similarities between the larval and adult *Drosophila* central nervous system. Elife 5, e13399.

Laughlin, S.B., de Ruyter van Steveninck, R.R., and Anderson, J.C. (1998). The metabolic cost of neural information. Nat Neurosci 1, 36–41.

Leader, D.P., Krause, S.A., Pandit, A., Davies, S.A., and Dow, J.A.T. (2018). FlyAtlas 2: a new version of the *Drosophila melanogaster* expression atlas with RNA-Seq, miRNA-Seq and sex-specific data. Nucleic Acids Res 46, D809–D815.

Lee, G., and Hall, J.C. (2001). Abnormalities of male-specific FRU protein and serotonin expression in the CNS of *fruitless* mutants in *Drosophila*. J Neurosci 21, 513–526.

Lee, G., Villella, A., Taylor, B.J., and Hall, J.C. (2001). New reproductive anomalies in *fruitless*-mutant *Drosophila* males: extreme lengthening of mating durations and infertility correlated with defective serotonergic innervation of reproductive organs. J Neurobiol 47, 121–149.

Lee, K.M., Daubnerova, I., Isaac, R.E., Zhang, C., Choi, S., Chung, J., and Kim, Y.J. (2015). A neuronal pathway that controls sperm ejection and storage in female *Drosophila*. Curr Biol 25, 790–797.

Lee, P.T., Zirin, J., Kanca, O., Lin, W.W., Schulze, K.L., Li-Kroeger, D., Tao, R., Devereaux, C., Hu, Y., Chung, V., et al. (2018). A gene-specific T2A-GAL4 library for *Drosophila*. Elife 7.

Li, H., Horns, F., Wu, B., Xie, Q., Li, J., Li, T., Luginbuhl, D.J., Quake, S.R., and Luo, L. (2017). Classifying *Drosophila* Olfactory Projection Neuron Subtypes by Single-Cell RNA Sequencing. Cell 171, 1206–1220 e1222.

Limmer, S., Weiler, A., Volkenhoff, A., Babatz, F., and Klambt, C. (2014). The *Drosophila* blood-brain barrier: development and function of a glial endothelium. Front Neurosci 8, 365.

Littleton, J.T., and Ganetzky, B. (2000). Ion channels and synaptic organization: analysis of the *Drosophila* genome. Neuron 26, 35–43.

Macosko, E.Z., Basu, A., Satija, R., Nemesh, J., Shekhar, K., Goldman, M., Tirosh, I., Bialas, A.R., Kamitaki, N., Martersteck, E.M., et al. (2015). Highly Parallel Genome-wide Expression Profiling of Individual Cells Using Nanoliter Droplets. Cell 161, 1202–1214.

Mamiya, A., Gurung, P., and Tuthill, J.C. (2018). Neural Coding of Leg Proprioception in *Drosophila*. Neuron 100, 636–650 e636.

Martin, C.A., and Krantz, D.E. (2014). *Drosophila melanogaster* as a genetic model system to study neurotransmitter transporters. Neurochem Int 73, 71–88.

Maurange, C., Cheng, L., and Gould, A.P. (2008). Temporal transcription factors and their targets schedule the end of neural proliferation in *Drosophila*. Cell 133, 891–902.

McGinnis, W., and Krumlauf, R. (1992). Homeobox genes and axial patterning. Cell 68, 283–302.

McIlroy, G., Foldi, I., Aurikko, J., Wentzell, J.S., Lim, M.A., Fenton, J.C., Gay, N.J., and Hidalgo, A. (2013). Toll-6 and Toll-7 function as neurotrophin receptors in the *Drosophila melanogaster* CNS. Nat Neurosci 16, 1248–1256.

Mendes, C.S., Bartos, I., Akay, T., Marka, S., and Mann, R.S. (2013). Quantification of gait parameters in freely walking wild type and sensory deprived *Drosophila melanogaster*. Elife 2, e00231.

Mendes, C.S., Rajendren, S.V., Bartos, I., Marka, S., and Mann, R.S. (2014). Kinematic responses to changes in walking orientation and gravitational load in *Drosophila melanogaster*. PLoS One 9, e109204.

Monastirioti, M., Linn, C.E., Jr., and White, K. (1996). Characterization of *Drosophila* tyramine beta-hydroxylase gene and isolation of mutant flies lacking octopamine. J Neurosci 16, 3900–3911.

Namiki, S., Dickinson, M.H., Wong, A.M., Korff, W., and Card, G.M. (2018). The functional organization of descending sensory-motor pathways in *Drosophila*. Elife 7.

Nässel, D.R. (2018). Substrates for Neuronal Cotransmission With Neuropeptides and Small Molecule Neurotransmitters in *Drosophila*. Front Cell Neurosci 12, 83.

Nässel, D.R., Holmqvist, M.H., Hardie, R.C., Hakanson, R., and Sundler, F. (1988). Histamine-like immunoreactivity in photoreceptors of the compound eyes and ocelli of the flies *Calliphora erythrocephala* and *Musca domestica*. Cell Tissue Res 253, 639–646.

Nässel, D.R., Pirvola, U., and Panula, P. (1990). Histaminelike immunoreactive neurons innervating putative neurohaemal areas and central neuropil in the thoraco-abdominal ganglia of the flies *Drosophila* and *Calliphora*. J Comp Neurol 297, 525–536.

Nässel, D.R., and Zandawala, M. (2019). Recent advances in neuropeptide signaling in *Drosophila*, from genes to physiology and behavior. Prog Neurobiol 179, 101607.

Niven, J.E., Graham, C.M., and Burrows, M. (2008). Diversity and evolution of the insect ventral nerve cord. Annu Rev Entomol 53, 253–271.

Noordermeer, J.N., Kopczynski, C.C., Fetter, R.D., Bland, K.S., Chen, W.Y., and Goodman, C.S. (1998). Wrapper, a novel member of the Ig superfamily, is expressed by midline glia and is required for them to ensheath commissural axons in *Drosophila*. Neuron 21, 991–1001.

Nusbaum, M.P., Blitz, D.M., and Marder, E. (2017). Functional consequences of neuropeptide and small-molecule co-transmission. Nat Rev Neurosci 18, 389–403.

Overend, G., Luo, Y., Henderson, L., Douglas, A.E., Davies, S.A., and Dow, J.A. (2016). Molecular mechanism and functional significance of acid generation in the *Drosophila* midgut. Sci Rep 6, 27242.

Özkan, E., Carrillo, R.A., Eastman, C.L., Weiszmann, R., Waghray, D., Johnson, K.G., Zinn, K., Celniker, S.E., and Garcia, K.C. (2013). An extracellular interactome of immunoglobulin and LRR proteins reveals receptor-ligand networks. Cell 154, 228–239.

Pan, Y., Robinett, C.C., and Baker, B.S. (2011). Turning males on: activation of male courtship behavior in *Drosophila melanogaster*. PLoS One 6, e21144.

Park, D., Veenstra, J.A., Park, J.H., and Taghert, P.H. (2008). Mapping peptidergic cells in *Drosophila*: where DIMM fits in. PLoS One 3, e1896.

Pauls, D., Blechschmidt, C., Frantzmann, F., El Jundi, B., and Selcho, M. (2018). A comprehensive anatomical map of the peripheral octopaminergic/tyraminergic system of *Drosophila melanogaster*. Sci Rep 8, 15314.

Pavlou, H.J., Lin, A.C., Neville, M.C., Nojima, T., Diao, F., Chen, B.E., White, B.H., and Goodwin, S.F. (2016). Neural circuitry coordinating male copulation. Elife 5.

Philippidou, P., and Dasen, J.S. (2013). Hox genes: choreographers in neural development, architects of circuit organization. Neuron 80, 12–34.

Pipes, G.C., Lin, Q., Riley, S.E., and Goodman, C.S. (2001). The Beat generation: a multigene family encoding IgSF proteins related to the Beat axon guidance molecule in *Drosophila*. Development 128, 4545–4552.

Prokop, A., and Technau, G.M. (1991). The origin of postembryonic neuroblasts in the ventral nerve cord of *Drosophila melanogaster*. Development 111, 79–88.

Rezával, C., Nojima, T., Neville, M.C., Lin, A.C., and Goodwin, S.F. (2014). Sexually dimorphic octopaminergic neurons modulate female postmating behaviors in *Drosophila*. Curr Biol 24, 725–730.

Rideout, E.J., Dornan, A.J., Neville, M.C., Eadie, S., and Goodwin, S.F. (2010). Control of sexual differentiation and behavior by the *doublesex* gene in *Drosophila melanogaster*. Nat Neurosci 13, 458–466.

Ro, J., Pak, G., Malec, P.A., Lyu, Y., Allison, D.B., Kennedy, R.T., and Pletcher, S.D. (2016). Serotonin signaling mediates protein valuation and aging. Elife 5.

Robinow, S., Campos, A.R., Yao, K.M., and White, K. (1988). The elav gene product of *Drosophila*, required in neurons, has three RNP consensus motifs. Science 242, 1570–1572.

Satija, R., Farrell, J.A., Gennert, D., Schier, A.F., and Regev, A. (2015). Spatial reconstruction of single-cell gene expression data. Nat Biotechnol 33, 495–502.

Schmid, A., Chiba, A., and Doe, C.Q. (1999). Clonal analysis of *Drosophila* embryonic neuroblasts: neural cell types, axon projections and muscle targets. Development 126, 4653–4689.

Schutzler, N., Girwert, C., Hugli, I., Mohana, G., Roignant, J.Y., Ryglewski, S., and Duch, C. (2019). Tyramine action on motoneuron excitability and adaptable tyramine/octopamine ratios adjust *Drosophila* locomotion to nutritional state. Proc Natl Acad Sci U S A 116, 3805–3810.

Seeds, A.M., Ravbar, P., Chung, P., Hampel, S., Midgley, F.M., Jr., Mensh, B.D., and Simpson, J.H. (2014). A suppression hierarchy among competing motor programs drives sequential grooming in *Drosophila*. Elife 3, e02951.

Sellami, A., Agricola, H.J., and Veenstra, J.A. (2011). Neuroendocrine cells in *Drosophila melanogaster* producing GPA2/GPB5, a hormone with homology to LH, FSH and TSH. Gen Comp Endocrinol 170, 582–588.

Shepherd, D., Harris, R., Williams, D.W., and Truman, J.W. (2016). Postembryonic lineages of the *Drosophila* ventral nervous system: Neuroglian expression reveals the adult hemilineage associated fiber tracts in the adult thoracic neuromeres. J Comp Neurol 524, 2677–2695.

Shepherd, D., Sahota, V., Court, R., Williams, D.W., and Truman, J.W. (2019). Developmental organization of central neurons in the adult *Drosophila* ventral nervous system. J Comp Neurol 527, 2573–2598.

Shirangi, T.R., Stern, D.L., and Truman, J.W. (2013). Motor control of *Drosophila* courtship song. Cell Rep 5, 678–686.

Sievers, F., and Higgins, D.G. (2018). Clustal Omega for making accurate alignments of many protein sequences. Protein Sci 27, 135–145.

Smarandache-Wellmann, C.R. (2016). Arthropod neurons and nervous system. Curr Biol 26, R960–R965.

Stahl, B.A., Peco, E., Davla, S., Murakami, K., Caicedo Moreno, N.A., van Meyel, D.J., and Keene, A.C. (2018). The Taurine Transporter Eaat2 Functions in Ensheathing Glia to Modulate Sleep and Metabolic Rate. Curr Biol 28, 3700–3708 e3704.

Suska, A., Miguel-Aliaga, I., and Thor, S. (2011). Segment-specific generation of *Drosophila* Capability neuropeptide neurons by multi-faceted Hox cues. Dev Biol 353, 72–80.

Tayler, T.D., Pacheco, D.A., Hergarden, A.C., Murthy, M., and Anderson, D.J. (2012). A neuropeptide circuit that coordinates sperm transfer and copulation duration in *Drosophila*. Proc Natl Acad Sci U S A 109, 20697–20702.

Thor, S., Andersson, S.G., Tomlinson, A., and Thomas, J.B. (1999). A LIM-homeodomain combinatorial code for motor-neuron pathway selection. Nature 397, 76–80.

Truman, J.W., and Bate, M. (1988). Spatial and temporal patterns of neurogenesis in the central nervous system of Drosophila melanogaster. Dev Biol 125, 145–157.

Truman, J.W., Moats, W., Altman, J., Marin, E.C., and Williams, D.W. (2010). Role of Notch signaling in establishing the hemilineages of secondary neurons in *Drosophila melanogaster*. Development 137, 53–61.

Truman, J.W., Schuppe, H., Shepherd, D., and Williams, D.W. (2004). Developmental architecture of adult-specific lineages in the ventral CNS of *Drosophila*. Development 131, 5167–5184.

Tsubouchi, A., Yano, T., Yokoyama, T.K., Murtin, C., Otsuna, H., and Ito, K. (2017). Topological and modality-specific representation of somatosensory information in the fly brain. Science 358, 615–623.

Tuthill, J.C., and Wilson, R.I. (2016). Parallel Transformation of Tactile Signals in Central Circuits of *Drosophila*. Cell 164, 1046–1059.

van der Maaten, L. (2014). Accelerating t-SNE using Tree-Based Algorithms. J Mach Learn Res 15, 3221–3245.

Venkatasubramanian, L., and Mann, R.S. (2019). The development and assembly of the *Drosophila* adult ventral nerve cord. Curr Opin Neurobiol 56, 135–143.

Volkenhoff, A., Weiler, A., Letzel, M., Stehling, M., Klambt, C., and Schirmeier, S. (2015). Glial Glycolysis Is Essential for Neuronal Survival in *Drosophila*. Cell Metab 22, 437–447.

von Philipsborn, A.C., Liu, T., Yu, J.Y., Masser, C., Bidaye, S.S., and Dickson, B.J. (2011). Neuronal control of *Drosophila* courtship song. Neuron 69, 509–522.

Williams, M.J., Goergen, P., Rajendran, J., Klockars, A., Kasagiannis, A., Fredriksson, R., and Schioth, H.B. (2014). Regulation of aggression by obesity-linked genes TfAP-2 and Twz through octopamine signaling in *Drosophila*. Genetics 196, 349–362.

Witt, E., Benjamin, S., Svetec, N., and Zhao, L. (2019). Testis single-cell RNA-seq reveals the dynamics of de novo gene transcription and germline mutational bias in *Drosophila*. Elife 8.

Wosnitza, A., Bockemuhl, T., Dubbert, M., Scholz, H., and Buschges, A. (2013). Inter-leg coordination in the control of walking speed in *Drosophila*. J Exp Biol 216, 480–491.

Xiong, W.C., Okano, H., Patel, N.H., Blendy, J.A., and Montell, C. (1994). repo encodes a glial-specific homeo domain protein required in the *Drosophila* nervous system. Genes Dev 8, 981–994.

Yang, C.P., Samuels, T.J., Huang, Y., Yang, L., Ish-Horowicz, D., Davis, I., and Lee, T. (2017). Imp and Syp RNA-binding proteins govern decommissioning of *Drosophila* neural stem cells. Development 144, 3454–3464.

Yellman, C., Tao, H., He, B., and Hirsh, J. (1997). Conserved and sexually dimorphic behavioral responses to biogenic amines in decapitated *Drosophila*. Proc Natl Acad Sci U S A 94, 4131–4136.

Zappia, L., and Oshlack, A. (2018). Clustering trees: a visualization for evaluating clusterings at multiple resolutions. Gigascience 7.

Zhou, B., Williams, D.W., Altman, J., Riddiford, L.M., and Truman, J.W. (2009). Temporal patterns of broad isoform expression during the development of neuronal lineages in *Drosophila*. Neural Dev 4, 39.

